# Cell-type specificity of neuronal excitability and morphology in the central amygdala

**DOI:** 10.1101/832295

**Authors:** Anisha P. Adke, Aleisha Khan, Hye-Sook Ahn, Jordan J. Becker, Torri D. Wilson, Spring Valdivia, Yae K. Sugimura, Santiago Martinez Gonzalez, Yarimar Carrasquillo

## Abstract

Central amygdala (CeA) neurons expressing protein kinase C delta (PKCδ^+^) or Somatostatin (Som^+^) differentially modulate diverse behaviors. The underlying features supporting cell-type-specific function in the CeA, however, remain unknown. Using whole-cell patch-clamp electrophysiology in acute mouse brain slices and biocytin-based neuronal reconstructions, we demonstrate that neuronal morphology and relative excitability are two distinguishing features between Som^+^ and PKCδ^+^ CeLC neurons. Som^+^ neurons, for example, are more excitable, compact and with more complex dendritic arborizations than PKCδ^+^ neurons. Cell size, intrinsic membrane properties, and anatomical localization were further shown to correlate with cell-type-specific differences in excitability. Lastly, in the context of neuropathic pain, we show a shift in the excitability equilibrium between PKCδ^+^ and Som^+^ neurons, suggesting that imbalances in the relative output of these cells underlie maladaptive changes in behaviors. Together, our results identify fundamentally important distinguishing features of PKCδ^+^ and Som^+^ cells that support cell-type-specific function in the CeA.

## INTRODUCTION

The laterocapsular subdivision of the central nucleus of the amygdala (CeLC) has received increasing interest due to its widespread function in mediating innate, as well as learned, adaptive and maladaptive behaviors. Previous work has demonstrated, for example, that the CeLC is functionally diverse, modulating fear conditioning and aversion (Aggleton, 2000, Davis and Whalen, 2001), nociception (Zald, 2003, Neugebauer et al., 2004, Veinante et al., 2013), anxiety, and drug reward and relapse in rodent models (Gilpin et al., 2015, Venniro et al., 2017, Venniro et al., 2018), to name a few. This diverse span of function is mirrored by the genetic, physiological and morphological heterogeneity in central amygdala (CeA) neuron subtypes (Schiess et al., 1999, Martina et al., 1999, Janak and Tye, 2015).

Two genetically identified cell types, protein kinase C delta-expressing (PKCδ^+^) neurons and somatostatin-expressing (Som^+^) neurons, constitute most CeLC neurons and are largely non-overlapping (Kim et al., 2017, Li et al., 2013, Wilson et al., 2019). PKCδ^+^ and Som^+^ neurons in the CeA undergo differential plasticity in the context of behavioral output and they both have critical, but distinct, functions in the modulation of CeA-dependent behaviors, including fear conditioning and pain-related behaviors. The activity of PKCδ^+^ cells, for example, is reduced following exposure to a conditioned stimulus after fear conditioning (Haubensak et al., 2010) but increased following nerve injury (Wilson et al., 2019). In contrast, Som^+^ cells respond to a threat stimulus by increasing their activity (Yu et al., 2016) but their excitability is decreased following nerve injury (Wilson et al., 2019). Consistent with these correlational changes in excitability, both Som^+^ and PKCδ^+^ CeA neurons have been shown to differentially contribute to fear and pain-related behaviors (Haubensak et al., 2010, Li et al., 2013, Wilson et al., 2019, Yu et al., 2016). The underlying features that support cell-type-specific opposite functions of these genetically distinct CeA neurons, however, remain unclear.

Previous studies have demonstrated that the electrophysiological and morphological properties of CeLC neurons are highly heterogenous across different species (Dumont et al., 2002, Chieng et al., 2006, Schiess et al., 1999, Li and Sheets, 2018). Recent studies have further shown that CeLC neurons with different firing properties are topographically organized based on their projection targets (Li and Sheets, 2018), suggesting that heterogeneity of function within the CeLC might also be anatomically defined.

In the present study, we contributed to the growing body of knowledge about the CeLC by performing a characterization of the electrophysiological and morphological properties of PKCδ^+^ and Som^+^ neurons. Our overarching hypothesis is that these two genetically distinct populations of CeLC neurons are electrophysiologically and morphologically different. We used whole-cell patch-clamp electrophysiology in acute mouse brain slices in combination with biocytin-based morphological reconstructions to characterize and compare the passive and active membrane properties, as well as the evoked repetitive firing responses, single action potential waveforms and neuronal morphologies of these two subpopulations of neurons. We further evaluated whether membrane properties and excitability are dependent on the anatomical localization within the central amygdala, both at the subnuclei and rostro-caudal levels.

Finally, using a mouse model of neuropathic pain, we tested whether perturbations known to alter CeLC-dependent behavioral outputs would result in a shift in the relative excitability of these two CeLC cell types. Using this cell-type-specific approach, we demonstrated that PKCδ^+^ and Som^+^ neurons have distinct electrophysiological and morphological properties and that the differences in the excitability of these cells are occluded in the context of neuropathic pain. Our combined findings provide an essential foundation for understanding functional heterogeneity within the CeA.

## RESULTS

### Som^+^ cells are more excitable than PKCδ^+^ neurons

Previous studies have shown cell-type-specific alterations in the firing responses of PKCδ^+^ and Som^+^ neurons following fear conditioning or nerve injury (Wilson et al., 2019, Yu et al., 2016, Ciocchi et al., 2010), demonstrating that plasticity in the firing responses of these cells underlies changes in behavioral output. Whether the relative excitability of PKCδ^+^ and Som^+^ neurons is different at baseline conditions, however, has not been determined. To do this, we crossed *Prkcd*-Cre or *Sst*-Cre mice with an Ai9 reporter strain to obtain offspring that expressed the fluorescent protein tdTomato in cells expressing PKCδ or Somatostatin, respectively (**Figure 1A**). Using an acute brain slice preparation, we performed whole-cell patch-clamp recordings from a total of 124 visually identified fluorescent neurons in the CeLC, corresponding to PKCδ^+^ and Som^+^ cells and distributed across the rostro-caudal extension of the CeLC (**Figure 1B and Figure 1 – figure supplement 1**). Consistent with previous studies (Chieng et al., 2006, Schiess et al., 1999, Martina et al., 1999, Lopez de Armentia and Sah, 2004), our experiments revealed that CeLC neurons display heterogenous firing responses (**Figure 1C**). Thus, three discrete firing phenotypes are observed: spontaneously active (S), late-firing (LF), and regular-spiking (RS) neurons. Late-firing neurons are silent at rest, fire repetitively in response to a prolonged (500 ms) depolarizing current injection and have a substantial delay to firing action potentials, while regular-spiking cells are also silent at rest and fire repetitively in response to depolarizing current injections but have a much shorter onset to action potentials firing.

**Figure 1.**
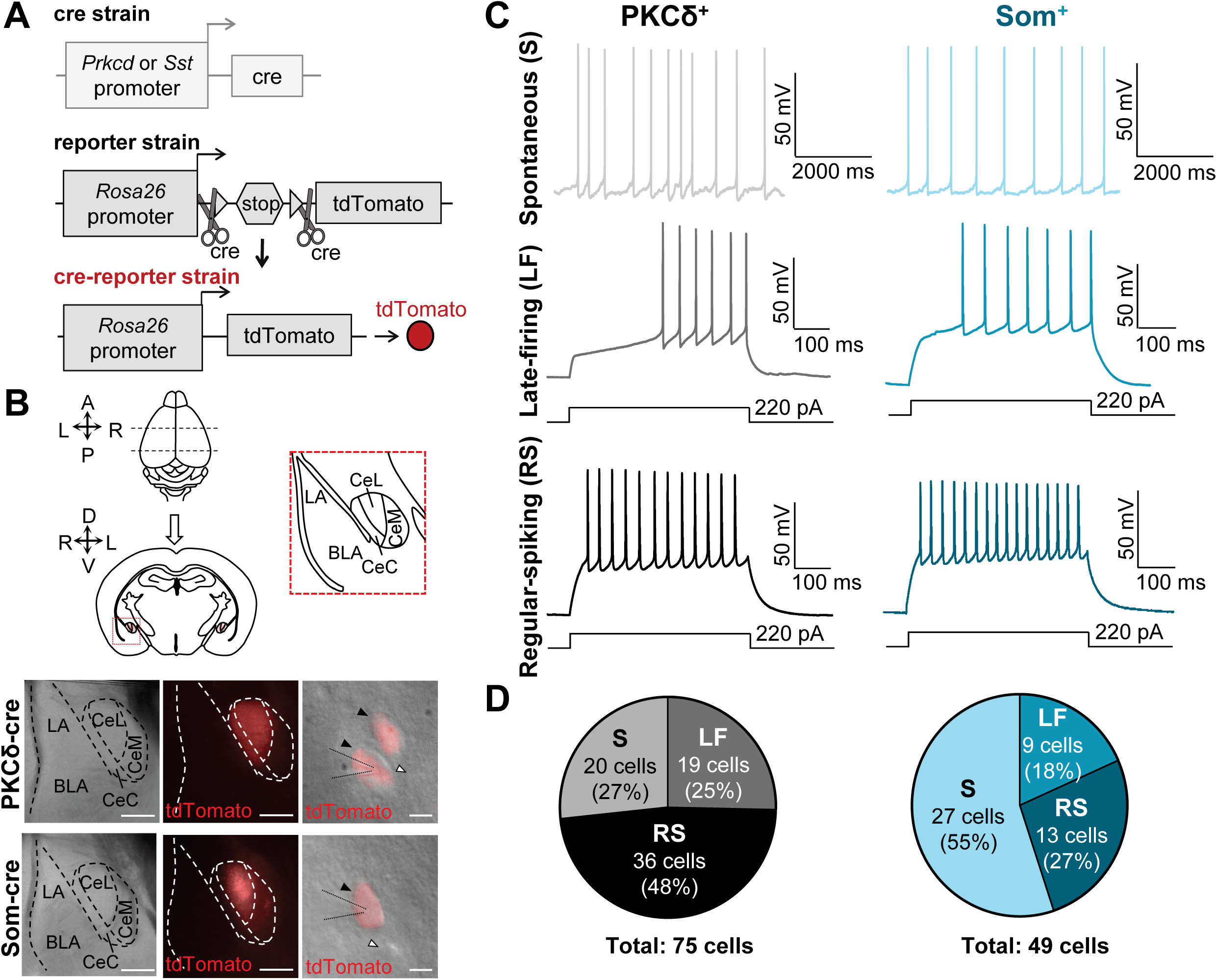
Firing phenotypes are heterogenous in PKCδ^+^ and Som^+^ CeLC neurons. (**A**) Strategy for labeling genetically-distinct subpopulations of neurons. *Pkcrd*-Cre or *Sst*-Cre mice were crossed with Ai9 reporter mice to produce offspring that express tdTomato fluorescent protein in either PKCδ^+^ or Som^+^ cells. (**B**) Acute amygdala slices for patch clamp electrophysiology. Whole brains were extracted and coronally sectioned. Bottom panels are low (left and middle) and high (right) magnification images of CeA slices. The CeA was visually identified by the distinct fiber bundles outlining the nuclei using differential interference contrast (left). PKCδ^+^ cells or Som^+^ cells expressing tdTomato (red) were readily seen under fluorescent microscopy (middle and right). Right panels show high magnification images of individual CeLC cells, with fluorescent images and differential interference contrast images overlaid. Black arrows denote fluorescently labeled cells, while white arrows denote unlabeled cells. Scale bars for left and center panel are 200 µm, right panel scale bar is 10 µm. (**C**) Representative voltage recordings of spontaneously active (S) cells, late-firing (LF) and regular-spiking (RS) PKCδ^+^ (left) or Som^+^ (right) neurons. (**D**) Proportions of each firing phenotype within recorded PKCδ^+^ and Som^+^ cell populations. The distribution of firing phenotypes is significantly (p = 0.0055, Chi-square test) different between PKCδ^+^ and Som^+^ cell populations. **See Figure 1 – figure supplement 1**.

As illustrated in the representative traces in **Figure 1C**, all three firing types are readily observed in both PKCδ^+^ and Som^+^ CeLC neurons. Quantification of the proportion of cells with different firing types revealed, however, that the distribution of firing phenotypes is significantly different between PKCδ^+^ and Som^+^ cells (**Figure 1D**). Of the 75 PKCδ^+^ cells recorded, for example, the majority (36/75; 48%) are regular-spiking neurons, whereas only 13 of the 49 (27%) Som^+^ neurons recorded are regular-spiking. In marked contrast, most (27/49; 55%) of the Som^+^ neurons recorded are spontaneously active at rest and only 20 of the 75 (27%) PKCδ^+^ neurons recorded are spontaneously active. The remaining PKCδ^+^ and Som^+^ cells were late-firing and the proportion of this firing type, relative to the total cells, is similar between PKCδ^+^ (19/75; 25%) and Som^+^ (9/49; 18%) neurons. Together, these results demonstrate that while firing phenotypes are heterogeneous in PKCδ^+^ and Som^+^ CeLC cells, the proportion of cells with different firing types is cell-type-specific. The greater proportion of spontaneously active Som^+^ cells suggests that these cells have a larger overall output compared to PKCδ^+^ cells in the CeLC.

The next set of experiments determined if the relative excitability of PKCδ^+^ and Som^+^ neurons is also different in late-firing and regular-spiking cells. Prolonged (500 ms) depolarizing current injections elicited repetitive firing in both late-firing and regular-spiking PKCδ^+^ and Som^+^ neurons, with firing responses increasing as a function of the current injection amplitude in all four cell types (**Figure 2A**). Evoked repetitive firing responses in late-firing and regular-spiking Som^+^ neurons are significantly higher than in late-firing and regular-spiking PKCδ^+^ cells, respectively, underscoring the notion that Som^+^ cell output far outpaces that of PKCδ^+^ neurons in the CeLC.

**Figure 2.**
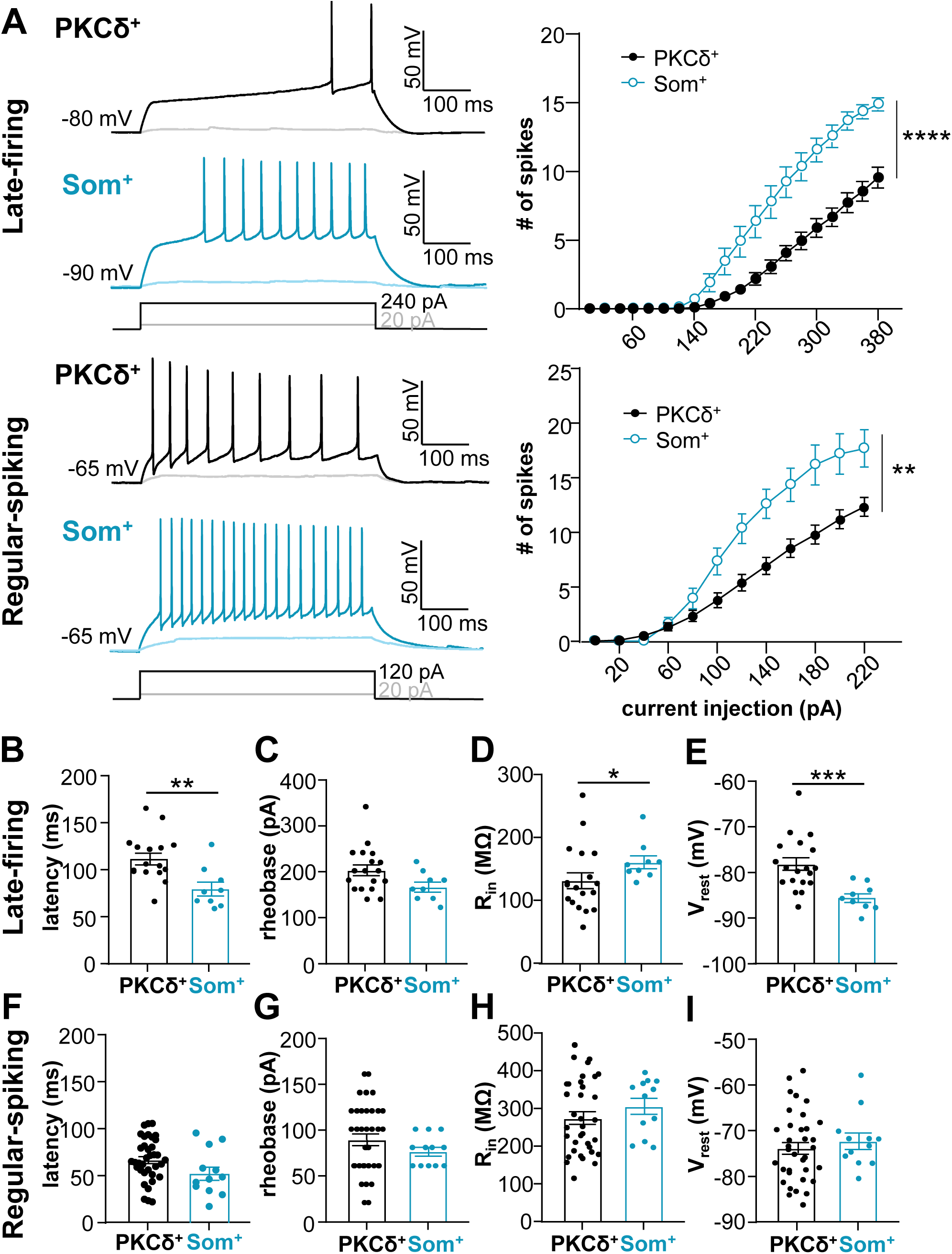
Som^+^ CeLC neurons are more excitable than PKCδ^+^ cells. (**A**) Representative voltage traces of late-firing (top left) or regular-spiking (bottom left) PKCδ^+^ cells (black) or Som^+^ cells (blue) in response to depolarizing current injections. Right panel shows the number of spikes elicited as a function of the current injection amplitude. ****p < 0.0001, **p < 0.0036, two-way ANOVA. (**B-F**) Latency to first spike (**B** and **F**), rheobase (**C** and **G**), input resistance (R_in_) (**D** and **H**), and resting membrane potential (V_rest_) (**E** and **I**) for late-firing (**B-E**) and regular-spiking (**F-I**) neurons. **p = 0.0039, unpaired two-tailed t-test; *p = 0.0308, Mann-Whitney U test; ***p = 0.0002, unpaired two-tailed t-test with Welch’s correction. For PKCδ^+^ cells: n = 16-19 cells for late-firing and n = 35-36 regular-spiking. For Som^+^ cells: n = 9 for late-firing and n = 12 for regular-spiking. All values are expressed as mean +/-S.E.M.

Consistent with the relative hyperexcitable phenotype of Som^+^ late-firing neurons, compared to PKCδ^+^ late-firing cells, the latency to first spike is significantly shorter in these cells, compared to PKCδ^+^ late-firing cells (**Figure 2B**). The minimal current amplitude that elicits an action potential (rheobase) as well as the voltage sag in response to a hyperpolarizing current injection, however, were indistinguishable between PKCδ^+^ and Som^+^ late-firing neurons (**Figure 2C and Figure 2 – figure supplement 1**).

Subthreshold membrane properties, like input resistance and resting membrane potential, can strongly influence the firing responses of a neuron in response to stimulation. To determine if subthreshold membrane properties contribute to the relative hyperexcitable phenotype of Som^+^ CeLC neurons, we measured and compared these two parameters in PKCδ^+^ and Som^+^ late-firing and regular-spiking neurons. Our analyses showed that input resistance is significantly higher in Som^+^, compared to PKCδ^+^ late-firing cells (**Figure 2D**), suggesting that differences in subthreshold conductances might contribute to the differences in excitability observed in PKCδ^+^ and Som^+^ neurons. Notably, the resting membrane potentials of late-firing Som^+^ neurons are significantly hyperpolarized relative to the resting membrane potentials in PKCδ^+^ late-firing cells (**Figure 2E**), demonstrating that the greater excitability of late-firing Som^+^ cells is independent of the resting membrane potential. In contrast, despite the pronounced differences in evoked firing responses of Som^+^ and PKCδ^+^ regular-spiking neurons, all other passive and active membrane properties measured are indistinguishable between these two genetically distinct cell types (**Figure 2F-I and Figure 2 – figure supplement 1**).

Together, these results demonstrate that the output of Som^+^ cells outpaces that of PKCδ^+^ cells in the CeLC but that the cellular mechanisms underlying the differences in excitability in Som^+^ and PKCδ^+^ neurons are distinct for late-firing and regular-spiking cells.

### PKCδ^+^, but not Som^+^, CeLC neurons display accommodation of repetitive firing

Inter-spike interval (ISI) accommodation reflects the ability of neurons to sustain the frequency of firing in response to prolonged depolarizing input. The presence of ISI accommodation has been previously reported in CeLC neurons and has been widely used as a parameter to classify neurons in this brain region (Schiess et al., 1999, Hunt et al., 2017, Zhu and Pan, 2004).

To further determine if firing properties in the CeLC are cell-type-specific and to gain additional insight into the mechanisms driving the differences in excitability of these cells, we measured and compared ISI accommodation between Som^+^ and PKCδ^+^ CeLC neurons. Prolonged (500 ms) current injections elicited repetitive firing in late-firing and regular-spiking Som^+^ and PKCδ^+^ CeLC neurons (**Figure 3A**). Quantification of the number of cells that display ISI accommodation further revealed that approximately half of the late-firing and half of the regular-spiking PKCδ^+^ neurons undergo ISI accommodation (**Figure 3B**). In marked contrast, however, only one of the 21 Som^+^ neurons analyzed exhibited ISI accommodation in response to depolarizing current injection. The almost complete lack of ISI accommodation in Som^+^ neurons is consistent with the overall higher output of these CeLC cells, compared to the PKCδ^+^ neurons. Further analyses of firing responses, as well as passive and active membrane properties, in accommodating and non-accommodating PKCδ^+^ neurons with either late-firing or regular-spiking phenotypes revealed that accommodating and non-accommodating neurons are mostly indistinguishable within firing types (**Table 1**). Thus, for the remainder of this study, data obtained from accommodating and non-accommodating PKCδ^+^ neurons were pooled and compared to non-accommodating Som^+^ cells.

**Figure 3.**
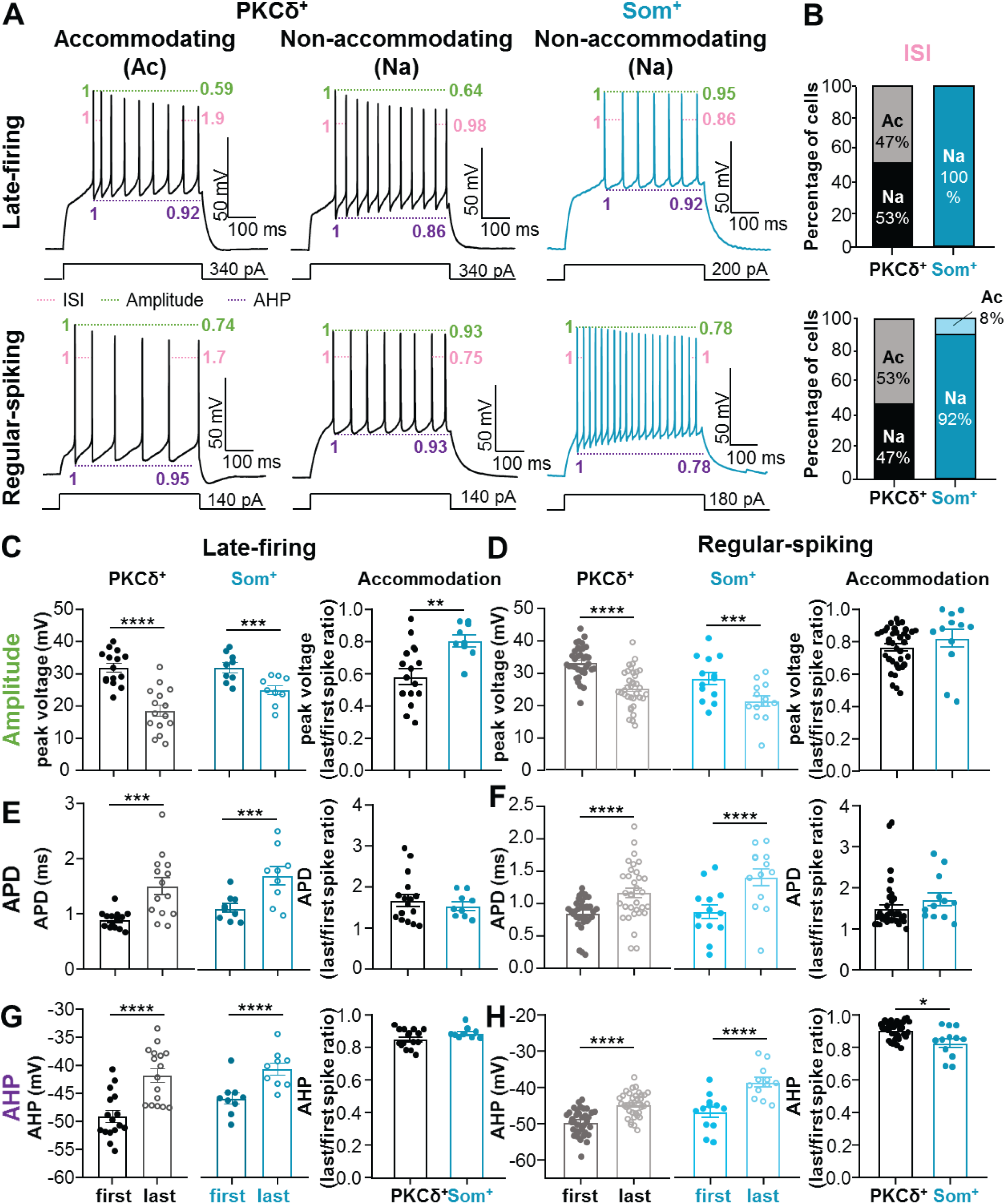
Accommodation is selective to PKCδ^+^, but not Som^+^, neurons. (**A**) Representative voltage records of PKCδ^+^ accommodating (Ac, left) and non-accommodating (Na, center) cells and Som^+^ Na cells (right) for late firing (top) regular spiking (bottom) cells. Pink annotations depict interspike interval (ISI) accommodation, green denotes spike amplitude accommodation, and purple shows afterhyperpolarization (AHP) amplitude accommodation. (**B**) The proportions of Na and Ac late-firing and regular-spiking PKCδ^+^ and Som^+^ cells. (**C-H**) Spike amplitude accommodation (**C-D**), action potential duration (APD) accommodation (**E-F**), and AHP amplitude accommodation (**G-H**) for late-firing (left) and regular-spiking (right) PKCδ^+^ and Som^+^ cells. ****p < 0.0001, ***p < 0.005, paired two-tailed t-test; **p = 0.0034, unpaired two-tailed t-test; *p = 0.0142, unpaired two-tailed t-test with Welch’s correction. For PKCδ^+^ cells: n = 14-16 cells for late-firing and n = 33-36 regular-spiking. For Som+ cells: n = 9 for late-firing and n = 12-13 for regular-spiking. All values are expressed as mean ± S.E.M.

**Table 1.**
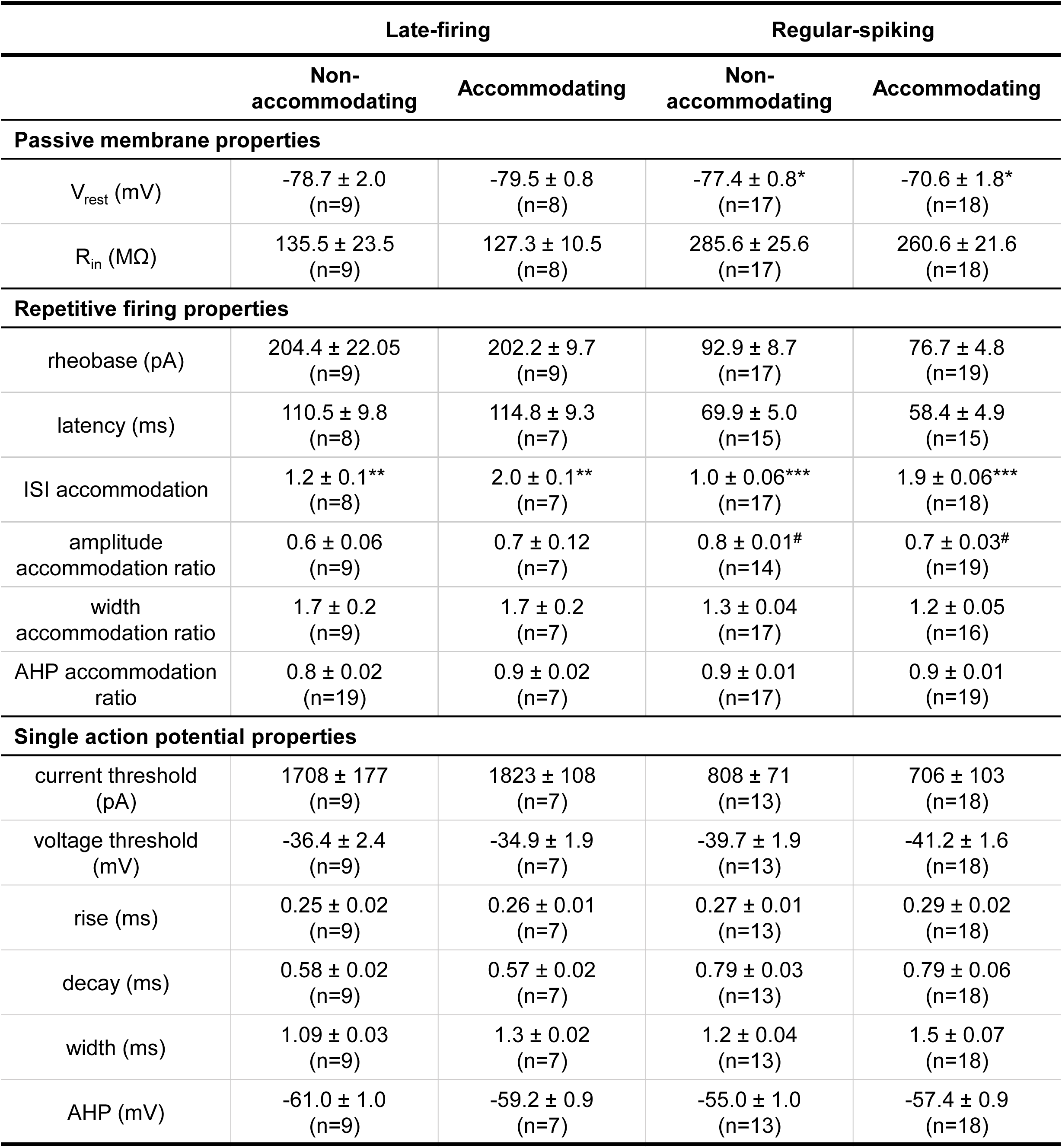
Passive membrane, repetitive firing, and single action potential properties of PKCδ^+^ non-accommodating and accommodating cells. All values are expressed as mean ± S.E.M. *p=0.0063, unpaired two-tailed t-test, comparing V_rest_ in regular-spiking accommodating and non-accommodating PKCδ^+^ cells. **p=0.0003, Mann Whitney U test, comparing ISI accommodation in accommodating and non-accommodating late-firing PKCδ^+^ cells; ***p<0.0001, unpaired two tailed t-test, comparing ISI accommodation in accommodating and non-accommodating regular-spiking PKCδ^+^ cells; ^#^p<0.0001, unpaired two-tailed t-test with Welch’s correction, comparing amplitude accommodation in regular-spiking PKCδ^+^ cells. V_rest_ = resting membrane potential; R_in_ = input resistance; ISI = inter-spike interval; AHP = afterhyperpolarization.

Frequency-dependent changes in spike amplitude, width and afterhyperpolarization (AHP) are three additional parameters used to measure the ability of neurons to sustain repetitive firing in response to prolonged depolarizing input. The presence of spike amplitude accommodation, spike broadening and AHP amplitude accommodation within an evoked train of action potentials are commonly used to classify and electrophysiologically characterize neurons in other brain regions, reflecting the repertoire of ion channels and ionic conductance of a cell (Bean, 2007). It is unknown, however, whether CeLC neurons display frequency-dependent changes in spike amplitude, width or AHP and, if they do, whether these changes are also cell-type-specific.

Measurements and comparisons of the amplitudes, widths and AHPs of the first and last spike within an evoked train of action potentials in late-firing and regular-spiking Som^+^ and PKCδ^+^ CeLC neurons demonstrated that all CeLC neurons display robust and significant frequency-dependent changes in spike amplitude, width and AHP amplitude in response to depolarizing current injections (**Figure 3C-H**). In all cells analyzed, for example, the amplitude of the last spike in an evoked train of action potentials is significantly shorter than the amplitude of the first spike within the same train (**Figure 3C-D**). Notably, while frequency-dependent shortening of the action potential is indistinguishable in PKCδ^+^ and Som^+^ regular-spiking neurons (**Figure 3D**), it is significantly larger in PKCδ^+^ than in Som^+^ late-firing cells (**Figure 3C**).

Quantification of action potential duration (APD) in the first and last spike further demonstrated that all four different cell types also exhibit robust APD accommodation, but that frequency-dependent APD broadening is indistinguishable between the different cell types (**Figure 3E-F**).

Finally, analysis of frequency-dependent AHP amplitude accommodation further revealed that all CeLC cell types exhibit significant AHP amplitude accommodation within an evoked train of action potentials and that AHP amplitude accommodation is significantly larger in Som^+^ regular-spiking than in PKCδ^+^ regular-spiking neurons (**Figure 3G-H**).

Our combined results, showing that ISI accommodation in response to depolarizing current injections is selective to PKCδ^+^ neurons and that frequency-dependent spike amplitude accommodation is more robust in PKCδ^+^ than in Som^+^, neurons demonstrates that the ability of PKCδ^+^ neurons to sustain firing in response to input is lower than that of Som^+^ cells and that differences in intrinsic membrane properties at the suprathreshold level contribute to these differences. These results are consistent with the findings discussed in the previous sections of this study and suggest that firing phenotypes in the CeLC are cell-type-specific, with Som^+^ cells displaying a larger overall output than PKCδ^+^ cells in the CeLC.

### Action potential repolarization is slower in Som^+^ than in PKCδ^+^ CeLC neurons

Suprathreshold membrane properties, including the membrane potential at which an action potential is initiated (voltage threshold) and the rates of depolarization and repolarization of individual action potentials, can also strongly influence neuronal excitability (Bean, 2007). To gain additional insight into the mechanisms underlying the differences in the excitability of PKCδ^+^ and Som^+^ cells, we examined the properties of single action potential waveforms elicited by a short (5 ms) depolarizing current injection in these cells (**Figure 4A**).

**Figure 4.**
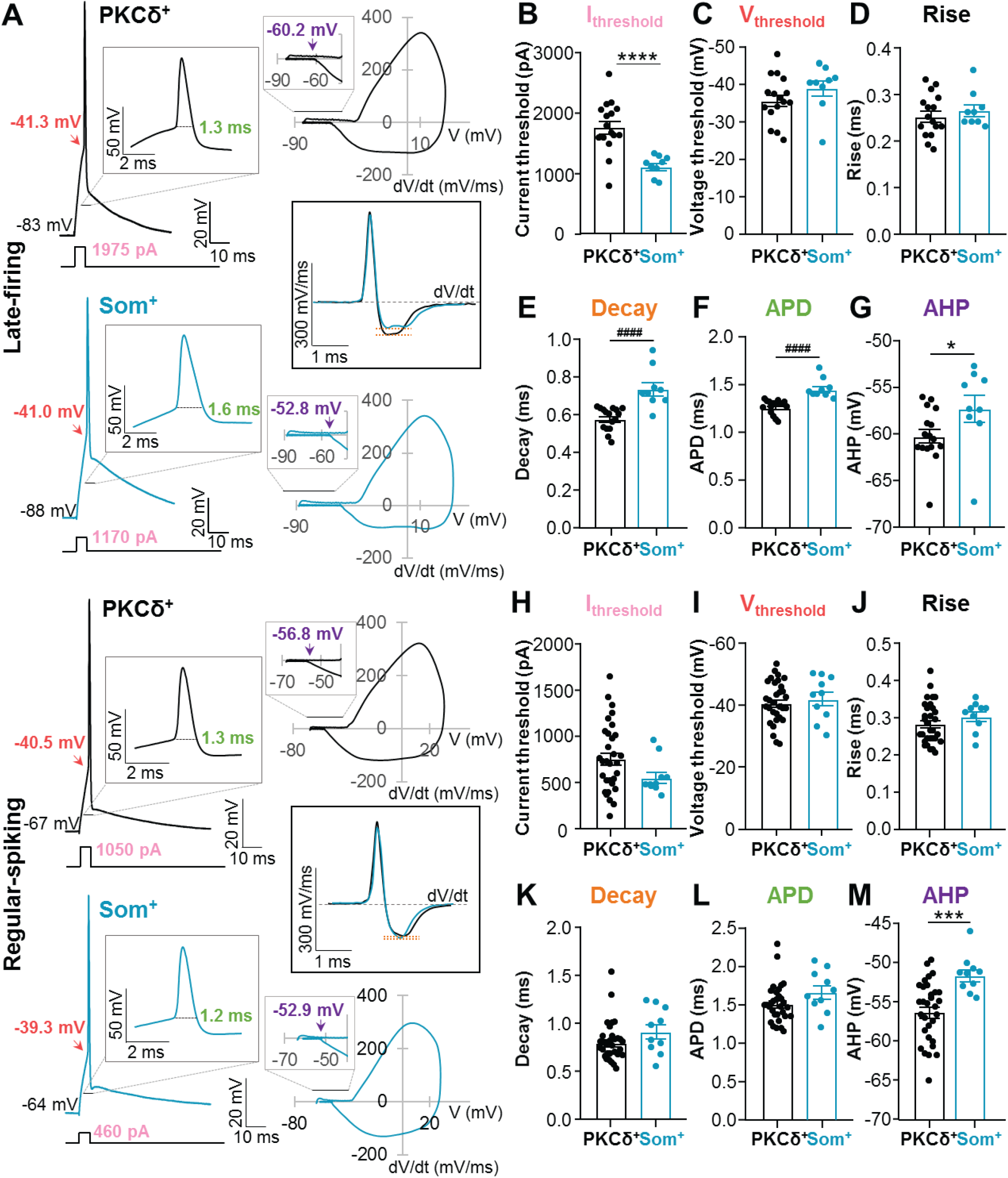
Slower repolarization in Som^+^, than in PKCδ^+^, neurons. (**A**) Representative single action potentials (left) elicited by 5 ms depolarizing current injections, phase plots (right) and plots of the first derivatives as a function of time (middle) of late-firing (top) and regular-spiking PKCδ^+^ (black) and Som^+^ (blue) neurons. Insets depict expanded time-scales. Single action potential analyses for late-firing (**B-G**) and regular-spiking (**H-M**) PKCδ^+^ and Som^+^ neurons. Current (**B** and **H**) and voltage (**C** and **I**) thresholds to fire a single action potential. Action potential rise time (**D** and **J**), action potential decay time (**E** and **K**), action potential duration (APD) (**F** and **L**) and aferhyperpolarization (AHP) amplitudes (**G** and **M**). ****p < 0.0001, unpaired two-tailed t-test with Welch’s correction; ^####^p < 0.0001, Mann-Whitney U test; *p = 0.0498, ***p = 0.0008, unpaired two-tailed t-test. For PKCδ^+^ cells: n = 16 cells for late-firing and n = 31 regular-spiking. For Som^+^ cells: n = 9 for late firing and n = 10 for regular spiking. All values are expressed as mean ± S.E.M.

Consistent with the hyperexcitable phenotype of Som^+^ late-firing neurons, the current amplitude required to induce an action potential (current threshold) is significantly smaller in Som^+^ than PKCδ^+^ late-firing neurons (**Figure 4B**). Analyses of the depolarizing phase of the action potentials further demonstrated that voltage thresholds and rise times are indistinguishable between PKCδ^+^ and Som^+^ late-firing cells (**Figure 4C-D**). In contrast, decay times are significantly longer, action potentials significantly prolonged and afterhyperpolarizations significantly depolarized in Som^+^, compared to PKCδ^+^ neurons (**Figure 4E-G**). These combined results demonstrate that while the depolarizing phase of action potentials is indistinguishable in PKCδ^+^ and Som^+^ late-firing CeLC neurons, the repolarizing phase is slower in Som^+^ than in PKCδ^+^ late-firing CeLC cells, likely contributing to the hyperexcitable phenotype observed in these cells.

Consistent with the indistinguishable subthreshold membrane properties and accommodation observed in PKCδ^+^ and Som^+^ regular-spiking CeLC neurons (**Figures 2 and 3**), most of the suprathreshold membrane properties measured are also indistinguishable in these cells (**Figure 4H-M**).

Together, these results suggest that differences in the intrinsic membrane properties of PKCδ^+^ and Som^+^ late-firing neurons contribute to the greater output of Som^+^ late-firing cells. The differences in excitability in Som^+^ and PKCδ^+^ regular-spiking neurons, however, seems to be independent of the intrinsic membrane properties of the cells, further supporting that the cellular mechanisms underlying the greater output of Som^+^ neurons are distinct for late-firing and regular-spiking CeLC cells.

### PKCδ^+^ neurons excitability is dependent on the rostro-caudal anatomical localization within the CeLC

Previous studies have shown that genetically distinct cells are differentially distributed throughout the CeLC (McCullough et al., 2018, Kim et al., 2017, Han et al., 2015, Wilson et al., 2019). As illustrated in **Figure 5**, and consistent with previous reports, for example, PKCδ^+^ cells are localized mostly to the lateral (CeL) and capsular (CeC) subdivisions of the CeLC, while Som^+^ cells are predominantly located in the CeL and medial subdivision (CeM) of the CeA (**Figure 5B**). Previous work has also demonstrated that although both cell types are found throughout the rostro-caudal axis, Som^+^ expression is greater in the anterior amygdala and decreases posteriorly, while PKCδ^+^ cells are expressed more abundantly in the medial CeLC (**Figure 5B**) (Han et al., 2015, Wilson et al., 2019).

**Figure 5.**
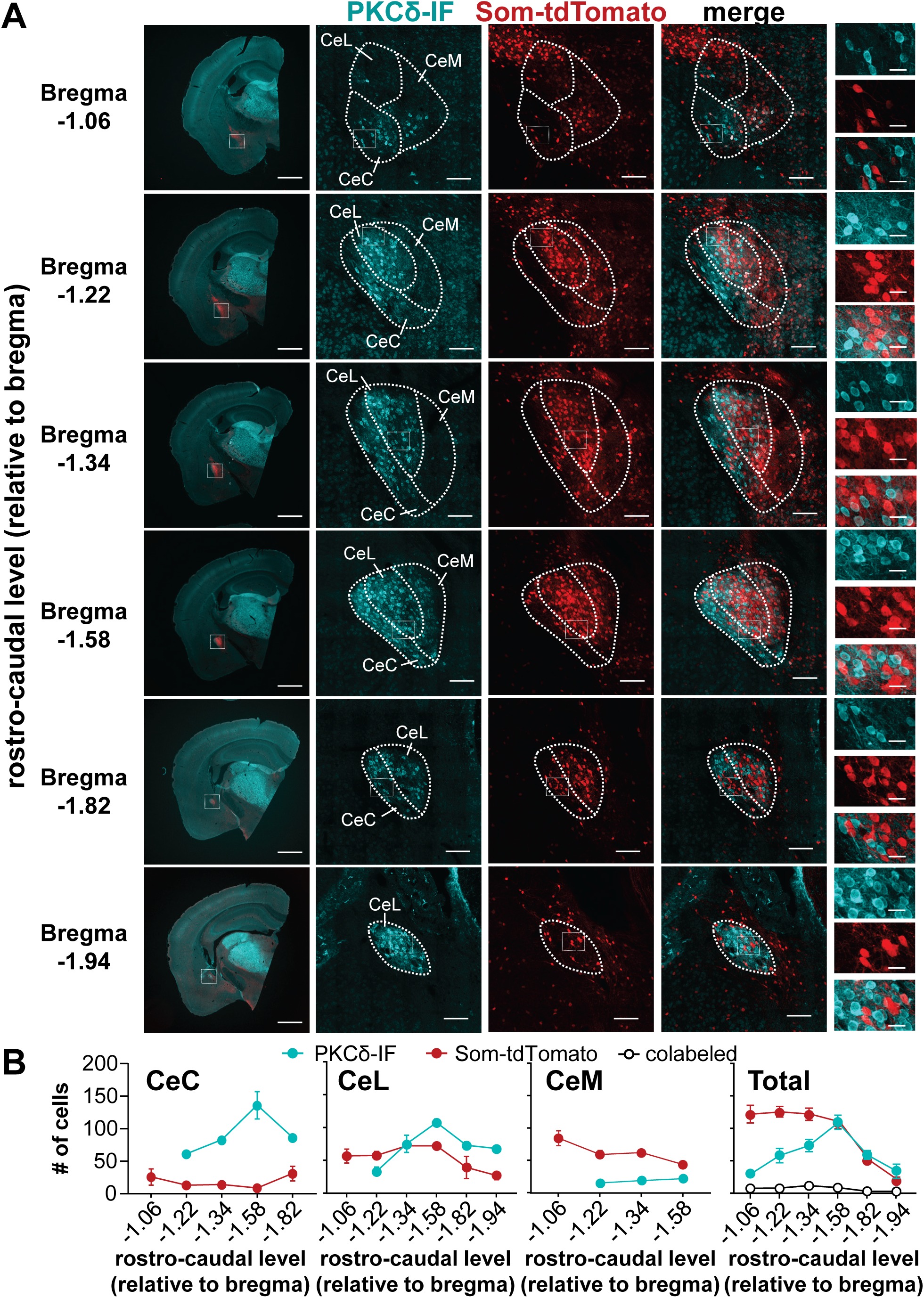
Rostro-caudal distribution of PKCδ^+^ and Som^+^ neurons in the CeA. (**A**) Representative low (left) and high (second to fifth panels) magnification images of coronal CeA slices immunostained for PKCδ (PKCδ-IF, cyan), with cells positive for Som-tdTomato shown in red. Merged signals between PKCδ-IF and Som-tdTomato are shown in the fourth panels. Rightmost panels depict high-magnification images of areas delineated by the white box. Scale bars are 1 mm for left panels, 100 µm for middle panels, and 20 µm for right panels. (**B**) Mean ± S.E.M. number of cells positive for PKCδ (cyan), Som (red) or colabeled with both (white circles) in the capsular (CeC), lateral (CeL) or medial (CeM) subdivisions of the CeA, as well as the total number of positive cells, are shown as a function of the rostro-caudal distribution relative to Bregma. n = 1-8 slices per rostro-caudal level from a total of 2-10 mice.

At the anatomical and electrophysiological levels, projection-specific neurons have been shown to be topographically organized within the CeLC and to exhibit distinct firing responses (Li and Sheets, 2018). It is unknown, however, whether the firing phenotypes of genetically distinct cells are dependent on their anatomical localization within the CeLC. We began to evaluate this by comparing the proportions of the three observed firing types (spontaneous, late-firing and regular-spiking; see **Figure 1**) in PKCδ^+^ and Som^+^ cells localized to different subnuclei or rostro-caudal levels within the CeLC.

Our analyses revealed that firing types of all PKCδ^+^ and Som^+^ cells are independent of their anatomical localization within the CeLC (**Table 2**). Further correlational analyses revealed, however, that the excitability of PKCδ^+^ regular-spiking neurons in the CeL correlates with the rostro-caudal localization of these cells (**Figure 6**). Thus, a significant positive correlation is seen for the number of spikes elicited in response to prolonged (500 ms) depolarizing current injection, with higher responses in neurons located in the posterior than in the anterior CeL (**Figure 6C**). Consistently, a significant negative correlation is observed for rheobase and latency to first spike, with lower values in neurons located in the posterior CeL, compared to neurons in the anterior CeL (**Figure 6D-E**). Firing responses to depolarizing current injections are indistinguishable in PKCδ^+^ regular-spiking neurons in the CeC as well as in all PKCδ^+^ late-firing neurons independently of their location within the rostro-caudal axis (**Figure 6C-E**).

**Figure 6.**
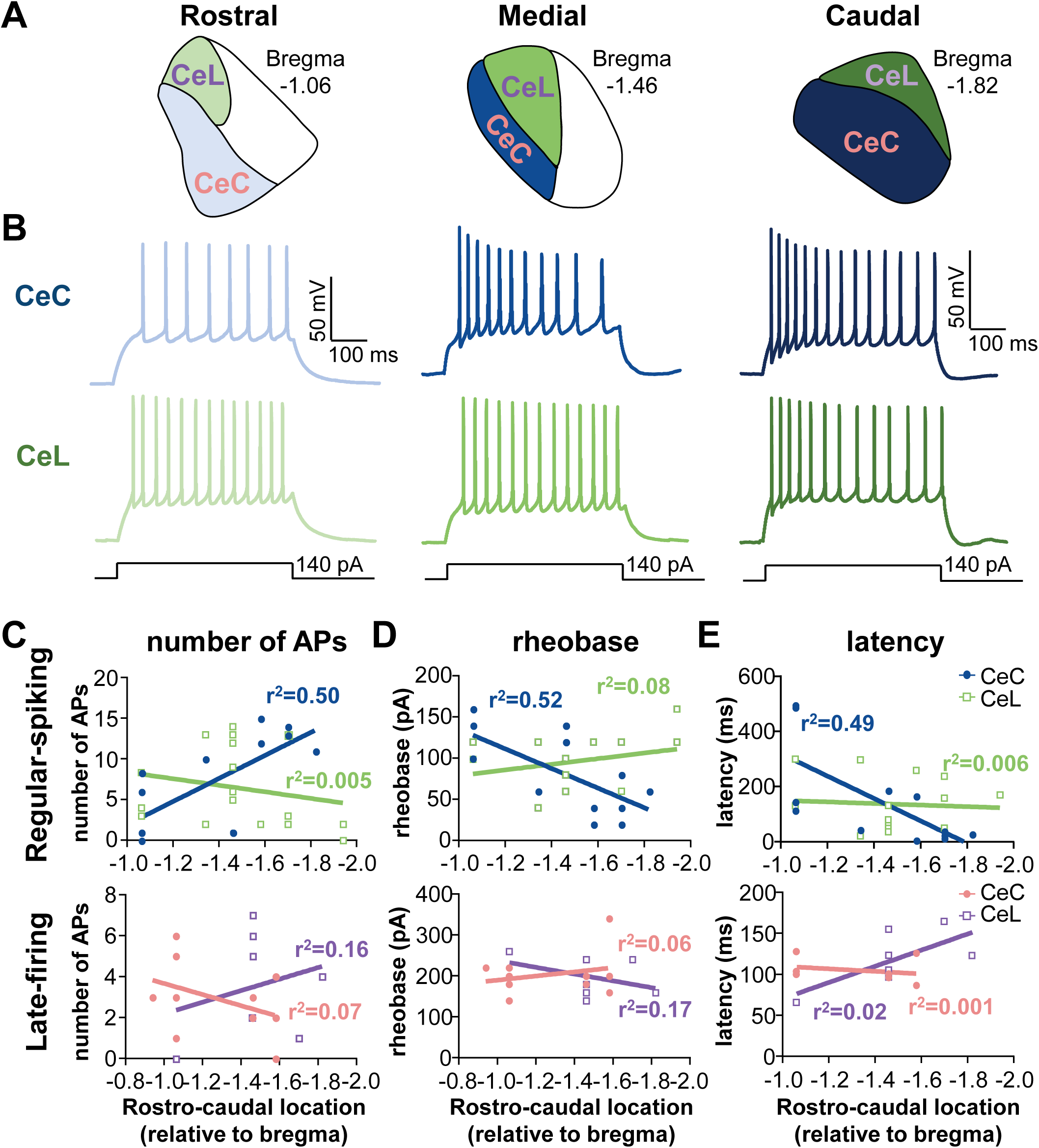
Firing responses in regular-spiking PKCδ^+^ neurons correlate with rostro-caudal anatomical location within the CeC. (**A**) Schematics of rostral, medial, and caudal regions of the CeLC, with the CeL represented in green and purple and the CeC represented in blue and pink. (**B**) Representative voltage traces of evoked firing responses in regular-spiking PKCδ^+^ neurons in the rostral, medial and caudal CeC (top panel, blue) and CeL (bottom panel, green). (**C-E**) Correlational plots between the number of evoked action potentials (**C**), rheobase (**D**), or latency to fire (**E**) and the rostro-caudal location of regular spiking (top) and late-firing (bottom) cells. Prolong (500 ms) depolarizing current injections of 140 pA and 240 pA were used to evoke repetitive firing in regular-spiking and late-firing cells, respectively. For regular-spiking neurons in the CeC, there was a positive correlation between the number of evoked action potentials and the rostro-caudal level (p = 0.0045, r^2^ = 0.5031, linear regression analysis) and a negative correlation between rheobase (p = 0.0036, r^2^ = 0.5193, linear regression analysis) and latency to first spike (p = 0.0118, r^2^ = 0.4858, linear regression analysis) with the rostro-caudal level. None of the measured parameters in the CeL and in late-firing cells in the CeC correlated with the rostro-caudal level. For CeL: n = 7 cells for late-firing and n = 18 for regular-spiking. For CeC: n = 10 for late-firing and n = 14 for regular-spiking.

**Table 2.** Firing phenotypes of PKCδ^+^ and Som^+^ CeLC cells by anatomical location. Proportions of cells with different firing phenotypes are shown for PKCδ^+^ and Som^+^ neurons in the capsular (CeC) and lateral (CeL) subdivisions of the central amygdala (CeA). Anterior is defined as the CeA between bregma -0.94 and bregma -1.34; medial as the CeA at bregma -1.46; and posterior as the CeA between bregma -1.58 and bregma -1.94. S = spontaneous; LF = late-firing; RS = regular-spiking; CeC = capsular subdivision of the central amygdala; CeL = lateral subdivision of the central amygdala.

|  | **Anterior**<br>(number of cells) | **Medial**<br>(number of cells) | **Posterior**<br>(number of cells) |
| --- | --- | --- | --- |
| **CeC-PKCδ<sup>+</sup>** |  |  |  |
| S | 4/14 | 0/4 | 2/11 |
| LF | 5/15 | 2/4 | 3/11 |
| RS | 6/15 | 2/4 | 6/11 |
| **CeL-PKCδ<sup>+</sup>** |  |  |  |
| S | 2/8 | 5/16 | 3/12 |
| LF | 1/8 | 4/16 | 3/12 |
| RS | 5/8 | 7/16 | 6/12 |
| **CeC-Som<sup>+</sup>** |  |  |  |
| S | 1/2 | 0/1 | 1/4 |
| LF | 0/2 | 0/1 | 2/4 |
| RS | 1/2 | 1/1 | 1/4 |
| **CeL-Som<sup>+</sup>** |  |  |  |
| S | 7/15 | 5/8 | 3/5 |
| LF | 4/15 | 1/8 | 1/5 |
| RS | 4/15 | 2/8 | 1/5 |

Together, these findings demonstrate that anatomical localization within the CeL is yet another source of heterogeneity that influences neuronal excitability in a cell-type specific manner in the CeA.

### PKCδ^+^ and Som^+^ neurons are morphologically distinct

It is widely known that neuronal morphology and dendritic spines impact the electrophysiological properties, and therefore cellular output, of neurons (Stiefel and Sejnowski, 2007, Mainen and Sejnowski, 1996, Connors and Regehr, 1996). While previous studies have demonstrated that neurons in the CeLC are both morphologically and electrophysiologically heterogeneous (Martina et al., 1999, Schiess et al., 1999, Chieng et al., 2006), a correlational link between the morphology and function of CeLC neurons is still missing.

Based on our electrophysiological findings demonstrating that excitability is markedly different in PKCδ^+^ and Som^+^ CeLC neurons, we hypothesized that these two subpopulations of CeLC cells are also morphologically distinct. To test this hypothesis, we filled some of the neurons that were used for the electrophysiological studies by including biocytin in the recording pipette solution (**Figure 7A**). A total of 7 PKCδ^+^ cells and 6 Som^+^ biocytin-filled cells were successfully recovered and reconstructed using this approach (**Figure 7B**).

**Figure 7.**
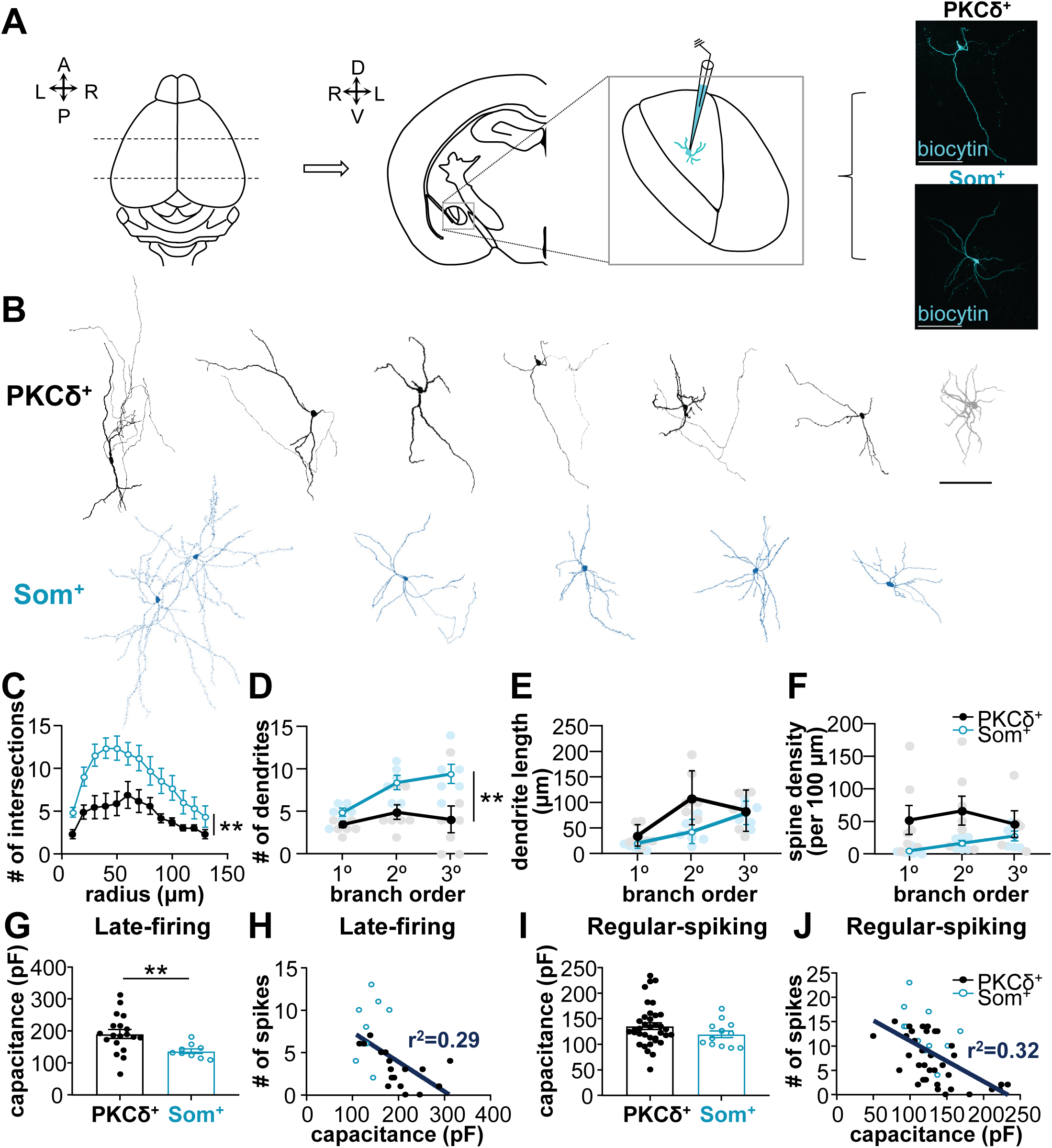
PKCδ^+^ and Som^+^ cells are morphologically distinct. (**A**) Morphological reconstruction of biocytin-filled cells in acute brain slices. CeLC cells were filled with biocytin during whole-cell patch-clamp recordings in acute amygdala slices. Representative images of biocytin-filled PKCδ^+^ and Som^+^ cells are shown in cyan in the right panel. Scale bars = 100 µm. (**B**) Morphological reconstruction of PKCδ^+^ (top) and Som^+^ biocytin-filled neurons. Scale bar = 100 µm. (**C**) Sholl analysis for number of dendritic intersections as a function of radial distance from soma. (**D-F**) Numbers (**D**), lengths (**E**), and spine densities (**F**) of primary (1°), secondary (2°), and tertiary (3°) dendrites for PKCδ^+^ and Som^+^ CeA cells. **p < 0.01, two-way ANOVA. (**G** and **I**) Whole-cell membrane capacitance for late-firing (**G**) and regular-spiking (**I**) PKCδ^+^ and Som^+^ CeA cells. (**H** and **J**) Correlational plots between the number of action potentials evoked in response to prolonged (500 ms) depolarizing current injections of either 140 (regular-spiking) or 240 pA (late-firing) and whole-cell membrane capacitance in late-firing (**H**) and regular-spiking (**J**) PKCδ^+^ (black) and Som^+^ (blue) CeA cells. A negative correlation was found in both late-firing (p = 0.0039, r^2^ = 0.29, linear regression analysis) and regular-spiking (p < 0.0001, r^2^ = 0.32, linear regression analysis) neurons. For PKCδ^+^ cells: n = 7 cells for morphology; n = 18-19 late-firing and n = 35-36 for regular-spiking. For Som^+^ cells: n = 6 cells for morphology; n = 9 late-firing and n = 13 for regular-spiking. All values are expressed as mean ± S.E.M.

Consistent with previous reports, the morphology of all CeLC neurons recovered resembled that of medium spiny neurons in the striatum. Notably, however, visual inspection of the reconstructed biocytin-filled neurons demonstrated that PKCδ^+^ cells display more polarity (triangular or bipolar) than Som^+^ cells, which have dendrites radiating in most directions outwards from the soma (**supplementary files 1 and 2**). One PKCδ^+^ cell (shown in grey in **Figure 7B**) was the only exception to this pattern. These qualitative observations suggested that the complexity of dendritic branching and dendritic length are different between PKCδ^+^ and Som^+^ CeLC neurons.

To quantify these qualitative differences in neuronal morphology, we performed Sholl analyses, which allows the quantification and comparison of the number of dendritic intersections as a function of distance from soma. The total number of primary, secondary and tertiary dendrites, as well as dendritic lengths and spine densities were also quantified in all cells.

As illustrated in **Figure 7C**, Sholl analyses revealed that the number of dendritic intersections is dependent on the distance from soma in both PKCδ^+^ and Som^+^ CeLC neurons, with maximal number of intersections observed at approximately 50 µm from the soma in both cell types. The number of dendritic intersections, however, was significantly higher in Som^+^ than in PKCδ^+^ neurons, demonstrating that dendritic arborizations are more complex in these cells compared to PKCδ^+^ cells. Consistent with the observed polarity of PKCδ^+^ cells, the total number of dendrites was significantly smaller in these neurons compared to the number of dendrites in Som^+^ cells (**Figure 7D**). In addition, posthoc analysis revealed that both the number and length of dendrites increases as a function of branch order in Som^+^ cells, but it is indistinguishable between primary, secondary, and tertiary dendrites of PKCδ^+^ cells (**Figure 7D-E**). Lastly, while dendritic spine densities increased as a function of branching order in Som^+^ neurons, it was indistinguishable between primary, secondary, and tertiary dendrites in PKCδ^+^ cells (**Figure 7F**).

Together, these results demonstrate that neuronal morphology differs in genetically distinct subpopulations of cells in the CeLC, with more complex dendritic branching patterns observed in Som^+^ neurons, than in PKCδ^+^ cells.

The combined results from our electrophysiological and morphological reconstruction of PKCδ^+^ and Som^+^ CeLC neurons strongly suggest that the morphological properties of PKCδ^+^ and Som^+^ neurons contribute to the differences in excitability displayed by these two populations of CeLC cells, with more compact Som^+^ neurons displaying higher excitability than the less compact PKCδ^+^ cells. To test this hypothesis, we used patch-clamp electrophysiology to measure and compare whole-cell capacitance in PKCδ^+^ and Som^+^ CeLC neurons. Whole-cell capacitance is commonly used to measure the total surface area of a cell, and therefore, reflects the size or compactness of a neuron, with lower whole-cell capacitance seen in smaller, more compact neurons and vice versa.

Consistent with the results of our morphological reconstructions that show Som^+^ neurons as more compact than PKCδ^+^ CeLC neurons, our electrophysiological measurements revealed that whole-cell capacitance is significantly lower in Som^+^ than in PKCδ^+^ CeLC late-firing neurons (**Figure 7G**). Notably, the number of evoked spikes significantly correlated with whole-cell capacitance in both late-firing and regular-spiking cells, with greater number of spikes seen in neurons with lower whole-cell capacitance (**Figure 7H and J**). These results demonstrate that more compact CeLC neurons are more excitable than larger cells, establishing a direct link between the distinct morphological properties of PKCδ^+^ and Som^+^ CeLC neurons and their excitability output.

### Nerve injury occludes differences in excitability between PKCδ^+^ and Som^+^ cells

The results of the experiments presented in **Figure 2**, performed in the absence of injury, show that Som^+^ cells in the CeLC are hyperexcitable compared to PKCδ^+^ neurons in this brain region. In a mouse model of neuropathic pain, however, previous studies have shown that nerve injury induces increases in the excitability of PKCδ^+^ neurons but that, in complete contrast, it decreases the excitability of Som^+^ CeLC cells (Wilson et al., 2019). These results suggest that nerve injury affects the excitability differences normally seen in PKCδ^+^ and Som^+^ neurons, ultimately affecting the overall output gain in the CeA. Whether and how cell-type-specific changes in excitability following nerve injury affect the relative output of PKCδ^+^ and Som^+^ neurons in the CeLC remains unknown.

To investigate this, we used the mouse cuff model of neuropathic pain in combination with whole-cell patch-clamp in acute brain slices (**Figure 8A**). Consistent with previous reports using this neuropathic pain model (Benbouzid et al., 2008, Wilson et al., 2019), cuff implantation in the sciatic nerve elicited robust and significant hypersensitivity to cold, heat and tactile stimulation in the hindpaw ipsilateral to treatment compared to the paw contralateral to cuff placement (**Figure 8A**). Cold, heat and tactile hypersensitivity were assessed using the acetone, Hargreaves and von-Frey tests, respectively.

**Figure 8.**
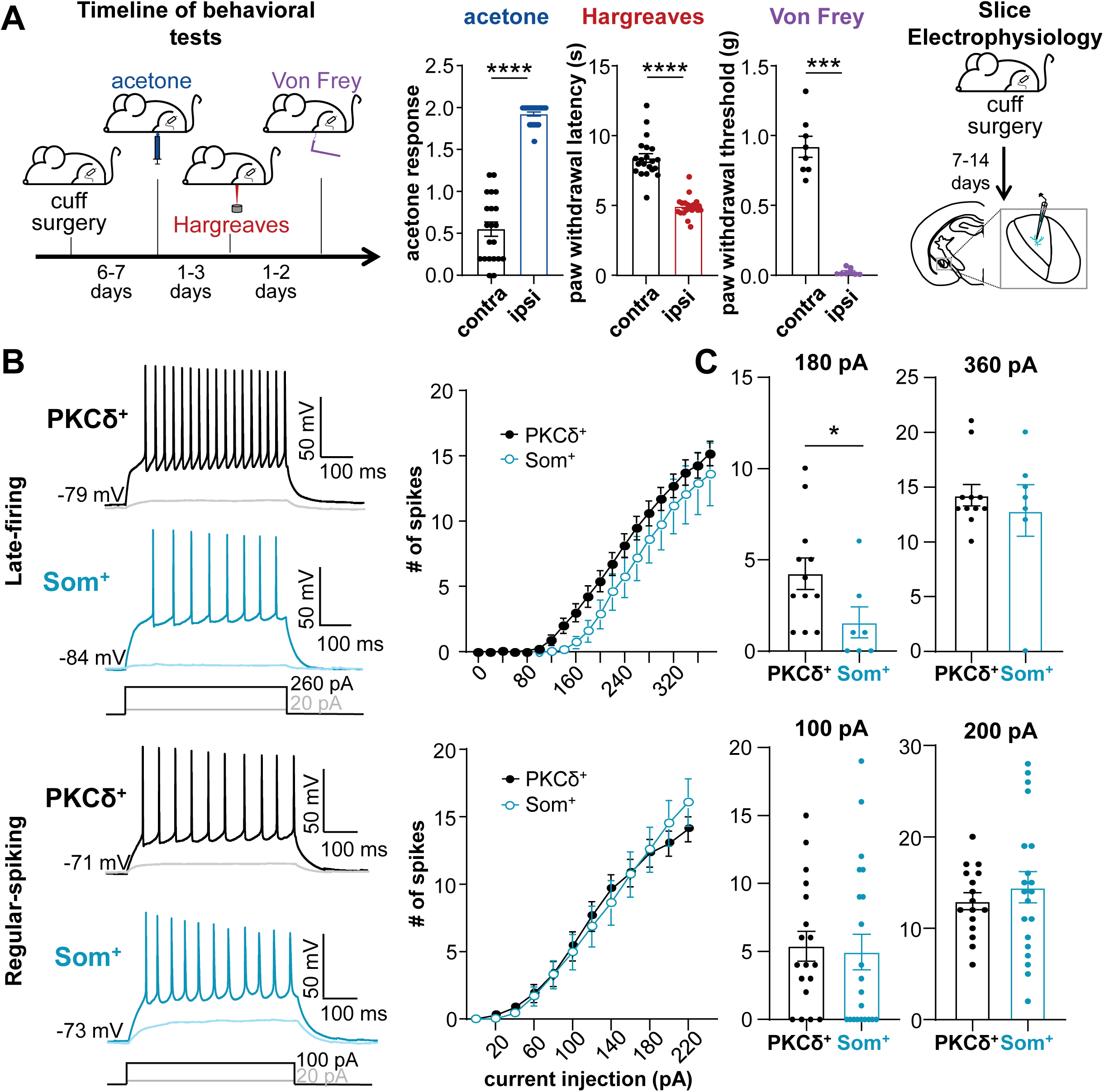
Excitability differences between PKCδ^+^ and Som^+^ cells are occluded in the context of persistent pathological pain. (**A**) Cuff model of neuropathic pain used in electrophysiological experiments. Following placement of the sciatic nerve cuff, mice developed hypersensitivity to cold (acetone test), heat (Hargreaves test), and tactile (von Frey test) stimulation on hindpaws contralateral to nerve injury, compared to the ipsilateral hindpaws. Acute brain slices for electrophysiological experiments were collected following cuff placement in the sciatic nerve of *Prkcd*-Cre::Ai9 or *Sst*-Cre::Ai9 mice. (**B**) Representative voltage recordings of late-firing (top) and regular-spiking (bottom) PKCδ^+^ (black) and Som^+^ (blue) cells in response to depolarizing current injections. Right panels show the number of spikes elicited as a function of the current injection amplitude. (**C**) The number of action potentials elicited in response to 180 pA and 360 pA depolarizing current injections in late-firing (top) and regular-spiking (bottom) PKCδ^+^ (black) and Som^+^ (blue) CeA cells. ****p < 0.0001, ***p < 0.0002, *p = 0.0314, Mann-Whitney U test. n = 8-21 mice for behavioral tests. For PKCδ^+^ cells: n = 11-12 late-firing and n = 16-18 for regular-spiking. For Som^+^ cells: n = 7 late-firing and n = 20-21 for regular-spiking. All values are expressed as mean ± S.E.M. **See Figure 8 – figure supplement 1**.

As illustrated in **Figure 8B**, and similar to what is seen in uninjured conditions (**Figure 2**), prolonged (500 ms) depolarizing current injections elicited repetitive firing in all four CeLC cell types following injury, with the number of evoked spikes increasing as a function of current injection amplitude. Notably, however, the number of spikes in response to prolonged depolarizing current injections is indistinguishable in PKCδ^+^ and Som^+^ neurons following injury, in both late-firing and regular-spiking cells. This is in marked contrast to the pronounced differences seen in uninjured animals (**Figure 2**) and demonstrates that nerve injury occludes differences in excitability between PKCδ^+^ and Som^+^ cells in the CeLC. Importantly, the differences in evoked firing responses between PKCδ^+^ and Som^+^ late-firing neurons is dependent on the amplitude of the depolarizing current injected (**Figure 8C**). Thus, while the number of spikes elicited by high amplitude (360 pA) current injection is indistinguishable in PKCδ^+^ and Som^+^ late-firing neurons, firing responses to low amplitude (180) current injections were significantly lower in Som^+^ than in PKCδ^+^ neurons in the CeLC. Firing responses of regular-spiking neurons, in contrast, were indistinguishable independently of the amplitude of current injected, supporting further that the mechanisms driving excitability of late-firing and regular-spiking neurons are distinct.

Consistent with the nerve injury-induced masking of neuronal excitability differences in PKCδ^+^ and Som^+^ CeLC neurons, the differences in resting membrane potential, input resistance, and latency to fire that we observed between PKCδ^+^ and Som^+^ CeLC late-firing neurons in uninjured conditions (**Figure 2B-E**) were also occluded in PKCδ^+^ and Som^+^ cells following nerve injury (**Figure 8 – figure supplement 1**).

Together, these results demonstrate that following nerve injury, the relative excitability of PKCδ^+^ and Som^+^ cells is disrupted, thus affecting the contribution of these cells to overall output in the CeLC.

## DISCUSSION

The CeLC has been recently hailed as a critical hub for modulating an array of behaviors, ranging from food-seeking to pain responses (Janak and Tye, 2015, Kim et al., 2017, Neugebauer et al., 2004). The two predominantly expressed cell types in the CeLC, PKCδ^+^ and Som^+^ cells, have been shown to modulate many of these behaviors, often in opposing ways (Wilson et al., 2019, Li et al., 2013, Ciocchi et al., 2010, Haubensak et al., 2010). In the present study, we show that PKCδ^+^ and Som^+^ CeLC neurons have different electrophysiological and morphological properties, supporting their distinct and diverse range of function. The results from our experiments demonstrate that while the firing phenotypes of these two genetically distinct CeLC cell types are heterogenous, there is a marked difference between the excitability of these cells, with Som^+^ neurons displaying a much greater output than PKCδ^+^ neurons.

In addition to the marked differences in excitability, our biocytin-based morphological reconstructions demonstrate that PKCδ^+^ and Som^+^ CeLC neurons are morphologically distinct, with more complex dendritic arborization patterns seen in Som^+^ than in PKCδ^+^ neurons. Importantly, our last set of experiments demonstrates that differences in the excitability of PKCδ^+^ and Som^+^ neurons are occluded in a mouse model of neuropathic pain, suggesting that maladaptive plastic changes that alter the relative output of CeLC cell types underlies the differential modulation of CeA-dependent behavior by these cells.

Together, the findings presented here identify fundamentally important differences in PKCδ^+^ and Som^+^ neurons that support the functional heterogeneity in the CeLC, shedding insight into how distinct subpopulations of neurons within this small brain structure can differentially contribute to the modulation of multiple behavioral outputs.

### Greater output is a common feature of all Som^+^ CeLC neurons

Previous studies have demonstrated that PKCδ^+^ and Som^+^ CeLC neurons have distinct, and often opposite, functions in the modulation of behaviors (Kim et al., 2017, Wilson et al., 2019, Janak and Tye, 2015). The cellular features that distinguish these two functionally distinct populations of CeLC neurons, however, are unknown. The results presented here demonstrate that PKCδ^+^ and Som^+^ neurons are electrophysiologically distinct. Despite the heterogeneity in firing responses in both cell types (**Figure 1**), a common and robust feature of all Som^+^ CeLC neurons is that they exhibit greater firing responses than PKCδ^+^ neurons within the same firing type (**Figure 2**). This is important because it suggests that the firing responses to input, as well as the overall output of these cells are distinct, demonstrating that information processing is different in PKCδ^+^ and Som^+^ cells at baseline. Differences in how these cells respond to input might, therefore, contribute to their selective or differential influence on behavioral outputs.

Identifying the source of cell-type-specific differential excitability in the CeLC is a crucial step towards understanding their opposite function. The results from the experiments presented here suggest that there are at least two distinct sources for the cell-type-specific differential excitability of PKCδ^+^ and Som^+^ CeLC cells. Differences in both passive and active intrinsic membrane properties, for example, seem to drive the relative hyperexcitability in Som^+^ late-firing neurons but do not contribute to the differences in excitability in regular-spiking cells (**Figures 2–4**). The greater input resistance, slower repolarization, shorter latencies and lower current thresholds for action potential generation, as well as the lack of ISI accommodation in late-firing Som+ neurons (compared to PKCδ^+^) are consistent with differences in potassium conductances between these cells.

Our results demonstrate that regular-spiking Som^+^ neurons are also much more excitable than PKCδ^+^ regular-spiking CeLC cells but, unlike the late-firing neurons, most of the passive and active membrane properties are indistinguishable in these cells (**Figures 2–4**). Since synaptic blockers were not used in our experiments, these results suggest that the higher output in regular-spiking Som^+^ neurons could be synaptically driven, which could result from higher excitatory inputs, lower inhibitory inputs, or a combination of both. The lateral and basolateral amygdala, as well as the lateral parabrachial nucleus are sources of excitatory inputs to the CeLC (Bernard and Besson, 1990, Lopez de Armentia and Sah, 2004). Previous work has demonstrated that both Som^+^ and Som^-^ CeLC cells receive monosynaptic excitatory inputs from these brain regions (Wilson et al., 2019, Li et al., 2013, Li and Sheets, 2019). In all these studies, however, glutamatergic inputs to Som^-^ cells are stronger than those to Som^+^ neurons in the CeC (Li and Sheets, 2019, Li et al., 2013). Since PKCδ^+^ and Som^+^ neurons comprise most of the CeLC and are mutually exclusive (Kim et al., 2017, Li et al., 2013, Wilson et al., 2019), it is likely that most of the Som^-^ neurons that receive stronger monosynaptic excitatory inputs are PKCδ^+^ cells. These results are somewhat counterintuitive because PKCδ^+^ cells show lower excitability than Som^+^ CeLC neurons, suggesting that differences in excitatory inputs do not contribute to higher excitability in Som^+^ CeLC neurons. Previous reports show, however, that PKCδ^+^ and Som^+^ CeLC neurons are interconnected and inhibit each other (Hunt et al., 2017, Haubensak et al., 2010), raising the possibility that differences in inhibitory inputs at the microcircuit level might contribute to the higher output of Som^+^ CeLC neurons. An important variable to consider when trying to integrate the results of the present study with the results of the synaptic studies mentioned above is that the experiments described here were conducted in the dark phase of the cycle whereas the synaptic experiments referenced above were performed in the light phase of the cycle. This is particularly important given recent work demonstrating that the sleep-wake state of an animal influences the activity of CeLC neurons (Ma et al., 2019). Defining the intrinsic and synaptic mechanisms underlying the differences in excitability between genetically distinct cells in the CeLC will be essential to fully understand their differential function in the modulation of behaviors.

### Cell-type-specific morphology as a predictor of function

CeLC neurons have been morphologically defined as medium spiny neurons with heterogenous dendritic branching complexities (Martina et al., 1999, Schiess et al., 1999, Chieng et al., 2006). Our biocytin-based morphological reconstruction of PKCδ^+^ and Som^+^ CeLC neurons revealed distinct morphological features in these cells (**Figure 7**). These results are surprising given the heterogenous firing phenotypes of these genetically distinct populations of cells (**Figure 1**) but, at the same time, are consistent with previous studies that have failed to correlate morphological properties of CeLC neurons with firing phenotypes (Amano et al., 2012, Chieng et al., 2006, Schiess et al., 1999).

The two common distinguishing features we found between Som^+^ and PKCδ^+^ CeLC neurons are their morphology and relative excitability. Thus, Som^+^ neurons are more compact, with lower whole-cell capacitance, a more complex dendritic branching pattern and a much greater overall firing output. PKCδ^+^ neurons, in contrast, have fewer (but longer) dendrites, higher whole-cell capacitance and a much lower overall firing output than Som^+^ neurons (**Figures 2 and 7 and supplementary files 1 and 2**).

Previous studies have shown similar correlations between morphology and excitability in striatal medium spiny and cortical pyramidal neurons (Gertler et al., 2008, van der Velden et al., 2012). Thus, neurons expressing the D2 dopamine receptor are compact, have lower whole-cell capacitance and are more excitable, (similar to our Som^+^ neurons) than those expressing the D1 receptor (Gertler et al., 2008). Similar to our findings in Som^+^ CeLC neurons, in the apical dendrite of layer 2/3 pyramidal neurons, higher dendritic branching complexities have been reported to correlate with greater excitability (van der Velden et al., 2012). Moreover, in cortical pyramidal neurons, higher complexity of dendritic branching complexity was shown to increase excitability by reducing ISI accommodation, which is consistent with the lack of ISI accommodation we see in Som^+^ neurons (**Figure 3**). Together, these results demonstrate that cell-type-specific morphology is an important determinant of neuronal excitability in PKCδ^+^ and Som^+^ CeLC neurons and can be used as a predictor of function in the CeA.

### Pain-related changes in excitability exemplify the ability of PKCδ^+^ and Som^+^ CeLC to undergo robust plasticity

Our cell-type-specific characterization of PKCδ^+^ and Som^+^ cells in the CeLC demonstrated that the overall output of these two genetically distinct populations is different at baseline (**Figure 2**). In the context of pain, however, we found that these differences were occluded (**Figure 8**), highlighting the power of these cells to undergo plasticity. Our findings suggest that a disruption in the excitability equilibrium of PKCδ^+^ and Som^+^ neurons can lead to CeLC-mediated pathological states. Whether the relative excitability of these cells is differentially affected in different behavioral contexts (i.e. food seeking behaviors, fear, drug reward and relapse, etc.) remains unknown.

Together, the findings described here demonstrate that genetically distinct CeLC neurons display cell-type-specific differences in firing output and dendritic morphology. These results support the distinct, and often opposite, contribution of PKCδ^+^ and Som^+^ CeLC neurons in the modulation of specific behavioral outputs and set the foundation for future studies aimed at identifying the cellular mechanisms driving heterogeneity of function in the CeA.

## MATERIALS AND METHODS

### Subjects

All animal procedures were performed in accordance with the guidelines of the National Institutes of Health (NIH) and were approved by the Animal Care and Use Committee of the National Institute of Neurological Disorders and Stroke and the National Institute on Deafness and other Communication Disorders. Adult (9- to 17-weeks-old) male mice were used for all experiments. *Prkcd*-cre heterozygote male or female mice (GENSAT-founder line 011559-UCD) were crossed with homozygous Ai9 mice (Jackson Laboratories). *Sst-*cre heterozygote males (Jackson Laboratory – founder line 018973) were crossed with homozygous female Ai9 (Jackson Laboratory) mice. Offspring mice were genotyped for the presence of cre-recombinase using DNA extracted from tail biopsies and PCR (Transnetyx) with the following primers: TTAATCCATATTGGCAGAACGAAAACG (forward) and CAGGCTAAGTGCCTTCTCTACA (reverse). Mice were housed in single cages or in pairs with littermates, separated by a perforated Plexiglass divider and kept in a reversed 12-hour light/dark cycle, with lights on from 9 pm to 9 am. Food and water were provided *ad libitum*. Prior to all experiments, mice were handled as previously described for at least 5 days to minimize potential stress effects associated with handling (Hurst and West, 2010). While handling, mice were also administered a 0.1 mL saline interperitoneally by the same experimenter that would be anesthetizing it for perfusion and acute slice preparation.

### *Ex-vivo* electrophysiology

#### Acute slice preparation

Mice were deeply anesthetized using 1.25% Avertin (0.4 mg/g body weight) injected intraperitoneally and then transcardially perfused with ice-cold cutting solution composed of (in mM) 110 choline chloride, 25 NaHCO_3_, 1.25 NaH_2_PO_4_, 2.5 KCl, 0.5 CaCl_2_, 7.2 MgCl_2_, 25 D-glucose, 12.7 L-ascorbic acid, and 3.1 pyruvic acid, oxygenated with 95%/5% O_2_/CO_2_. The brains were rapidly extracted, placed in ice-cold cutting solution, and cut in coronal slices (250 - 300 µm) using a Leica VT1200 S vibrating blade microtome (Leica Microsystems Inc., Buffalo Grove, IL, USA). Slices containing the CeA were incubated at 33°C for 30 minutes in a holding chamber containing artificial cerebral spinal fluid (ACSF) composed of (in mM) 125 NaCl, 2.5 KCl, 1.25 NaH_2_PO_4_, 25 NaHCO_3_, 2 CaCl_2_, 1 MgCl_2_, and 25 D-glucose. The chambers containing the slices were then moved to room temperature, and slices recovered for at least 20 minutes prior to recording. During incubation and recovery, the chambers were continuously oxygenated with 95%/5% O_2_/CO_2_.

#### Whole-cell patch-clamp recordings

The recording chamber was perfused continuously with ACSF oxygenated with 95%/5% O2/CO2 (1 mL/min) and all recordings were performed at 33±1°C. A recording chamber heater and an in-line solution heater (Warner Instruments) were used to control and monitor the bath temperature throughout the experiment. Recording pipettes (3-5 MΩ resistance) were filled with internal solution composed of (in mM) 120 potassium methyl sulfate, 20 KCl, 10 HEPES, 0.2 EGTA, 8 NaCl_2_, 4 Mg-ATP, 0.3 Tris-GTP, and 14 phosphocreatine with a pH of 7.3 using 5 M KOH and an osmolarity of ∼300 mosmol^-1^. Biocytin (3 mg/mL) was added to the internal solution of some recordings and sonicated in ice-cold water for 20 minutes. Whole-cell current-clamp recordings were obtained from tdTomato-expressing CeLC neurons. Cells were visually identified using an upright microscope (Nikon Eclipse FN1) equipped with differential interference contrast optics with infrared illumination and epifluorescence. Recording electrodes were visually positioned in the CeLC, guided by the distinctive fiber bundles and anatomical landmarks delineating its structure (**Figure 1B**). Recordings were controlled using the Multiclamp 700B patch-clamp amplifier interfaced with a Digidata 1500 acquisition system and pCLAMP 10.7 software (Axon Instruments, Union City, CA) on a Dell computer. Before forming a membrane-pipette seal, pipette tip potentials were zeroed and pipette capacitances and series resistances (not exceeding 20 MΩ) were monitored throughout the recordings. Whole-cell capacitance was measured in voltage-clamp configuration, with the cell held at -70 mV then subjected to a 25 ms ±10 mV current change. Brief (5 ms) and prolonged (500 ms) depolarizing current of various amplitudes were injected, while the cells were at resting membrane potential, to elicit single and repetitive action potential firing, respectively.

#### Data Analysis

Electrophysiological data were analyzed using ClampFit 10.7 (Molecular Devices), Microsoft Excel, Mini Analysis (v. 6.0.7, Synaptosoft, Inc., Decatur, GA), and Prism (v. 8, GraphPad Software Inc., La Jolla, CA). Data obtained from naïve animals and animals that received a sciatic nerve sham surgery were pooled and used to analyze baseline properties as no significant differences were seen between these conditions. Single action potential properties were measured from the action potentials generated in response to a 5 ms depolarizing current injection. Current threshold for action potential generation (I_threshold_) was defined as the minimum current injection required to elicit an action potential. Voltage threshold (V_threshold_) was calculated from the third derivative of the variation in membrane potential as a function of time during the rise of the action potential using the Mini Analysis software. Differentiated traces were digitally Gaussian filtered and smoothed by 30-100 points. Action potential duration (APD) was measured at 100% repolarization to voltage threshold (V_threshold_). Rise time was defined as the time required for the membrane potential to reach peak voltage from V_threshold_ and decay was defined as the time required for the membrane potential to repolarize from 90% of its peak to V_threshold_. Phase plots of single action potentials were generated by plotting the first derivative of the variation in membrane potential as a function of the membrane potential. Action potential afterhyperpolarization (AHP) was calculated from the phase plots of single action potentials and was defined as the voltage at which the first derivative of the variation in membrane potential during the repolarizing phase of the action potential reached zero or switched polarity. Input resistance (R_in_) was calculated using the average change in membrane potential in response to a ±20 pA current injection of 500 ms duration. Rheobase was defined as the minimum current required to induce an action potential in response to a 500 ms depolarizing current injection. Latency to fire for PKCδ^+^ vs Som^+^ cells comparisons was calculated at 2x rheobase and was defined as the time between current injection onset to action potential threshold. Voltage sag was calculated from the difference between the steady state and peak voltage responses to a 500 ms 500 pA hyperpolarizing current injection. Accommodation of inter-spike interval (ISI), action potential amplitude, action potential duration (calculated at 50% repolarization relative to voltage threshold) and afterhyperpolarization amplitude were calculated from the ratio of the measurements obtained from the last and first action potential in response to a 500 ms depolarizing current injection at 2x rheobase. ISI accommodating cells were defined as cells with a ratio greater than or equal to 1.5 whereas ISI non-accommodating cells had a ratio of less than 1.5. Latency to first spike was used to classify cells as late-firing or regular-spiking neurons. Cells with latencies shorter than 100 ms (at baseline) or 90 ms (pain conditions) were classified as regular-spiking. Conversely, cells with latencies higher than 100 ms (at baseline) or 90 ms (pain conditions) were classified as late-firing. Current amplitudes that elicited an average of 10 spikes (range of 5-19 spikes) were used to calculate latencies to first spike and, subsequently, to classify cells as late-firing or regular-spiking. In baseline (no pain) conditions, this current amplitude was 220 pA for Som^+^ cells and 280 pA for PKCδ^+^ neurons. Current amplitudes of 220 pA were used for both Som^+^ and PKCδ^+^ neurons in pain conditions. Number of spikes in response to a 500 ms depolarizing current injection of 140 pA (regular-spiking cells) or 240 pA (late-firing cells) amplitude was used to evaluate subnuclei and rostro-caudal differences in firing responses as well as firing responses as a function of whole-cell capacitance. Whole-cell membrane capacitance was calculated by integrating the capacitive transients elicited by a 25 ms voltage step (±10 mV) from -70 mV. Recording sites were constructed using the Mouse Brain Atlas as a guide (Paxinos et al., 2001).

### Morphological reconstruction of biocytin-filled cells

#### Preservation and staining of biocytin-filled neurons

Following current-clamp recordings, we followed procedures previously described to remove the recording electrode from the cell and retain the morphology of the cell (Swietek et al., 2016). In brief, the recording pipette was moved in slow alternating steps upward and outward in voltage-clamp mode while continuously monitoring the capacitive transients in order to reestablish a seal. Following re-sealing, the slice was left in the recording chamber for approximately three minutes to ensure transport of biocytin to distal processes. The slice was then removed from the recording chamber and immediately placed into 4% paraformaldehyde solution in 0.1 M Phosphate Buffer (PFA/PB), pH 7.4, at 4°C for 48 hours, followed by 0.1 M Phosphate Buffered Saline (PBS) (pH 7.4) with 0.01% sodium azide at 4°C until staining. Slices were rinsed with 0.1 M PBS 3 times for 5 minutes at room temperature while shaking at a low speed, then incubated in PBS containing 0.1% Triton-X-100 for 10 minutes. Samples were then incubated overnight, at 4°C and protected from light, in 1:500 Alexa Fluor 647 Streptavidin (Jackson 016-600-084) in blocking solution containing 1.5% normal goat serum (NGS) (Vector Labs, Burlingame, CA), 0.1% Triton-X-100, 0.05% Tween-20, and 1% bovine serum albumin (BSA). In minimal light, slices were then washed in 0.1 M PBS 4 times for 30 minutes at room temperature. Slices were then cleared using increasing concentrations of 2,2’-thiodiethanol (TDE) for 10 minutes each using 10, 30, 60, and 80% concentrations (Costantini et al., 2015), followed by incubation in 97% TDE for two hours. Slices were then mounted on positively-charged glass slides and covered with glass coverslips using 97% TDE.

#### Image acquisition, morphological reconstruction and analysis

Images of recovered biocytin-filled neurons were taken using a Nikon A1R laser scanning confocal microscope with a 40x oil-immersion objective. Gain and pinhole size were kept constant between experiments. Sequential acquisition of z-stacks was collected at 0.09 μm steps. Images were collected at varying sizes, depending on the extension of the dendrites of each neuron, and were then automatically stitched upon acquisition using NIS Elements software. To quantify neuronal morphology, z-stacks were analyzed using Neurolucida 360 (NeuroLucida 360, MBF Bioscience, Williston, VT). Cell bodies were identified using automatic detection of a user-defined area and dendritic branches were traced using the user-guided tree-tracing function. To identify spines, automatic spine detection was performed using image noise filtering. The experimenter manually confirmed or rejected all parameters analyzed. Primary dendrites were defined as the dendrites directly extending from the soma. Secondary dendrites branched from primary dendrites, and tertiary dendrites were the branching points of secondary dendrites. Dendritic length was defined as the distance from the trunk of the branch to either the distal branching point, or the visible end of the projection. Spine density was calculated by counting the individual spines present on the dendrites and reported per 100 µm of dendrite. Dendritic properties were obtained using branched-structure and Sholl analysis functions from NeuroLucida Explorer. The number of dendritic intersections at increasing radial distances from the soma, in 10 µm increments, were quantified and used for the Sholl analysis.

### Immunohistochemistry

Sst-cre::Ai9 mice were deeply anesthetized with 1.25% Avertin (2,2,2-tribromoethanol and tert-amyl alcohol in 0.9% NaCl; 0.025 ml/g body weight) and were then perfused transcardially with 37°C 0.9% NaCl, followed by 100 mL of ice-cold 4% PFA/PB. The brains were carefully dissected and post-fixed overnight in 4% PFA/PB at 4°C. After cryoprotection in 30% sucrose/PB for 48 h, the brains were sliced in 30 μm coronal sections using a freezing sliding microtome. The sections were stored in 0.1 M Phosphate Buffered Saline (PBS), pH 7.4 containing 0.01% sodium azide (Sigma) at 4°C until immunostaining. To begin staining, sections were rinsed in PBS, then incubated in PBS containing 0.1% Triton X100 for 10 minutes at room temperature, followed by blocking at room temperature for 30 minutes in blocking solution containing 5% normal goat serum (NGS) (Vector Labs, Burlingame, CA), 0.1% Triton X100, 0.05% Tween-20 and 1% bovine serum albumin (BSA). Sections were then incubated for 72 h at 4°C in mouse anti-PKCδ primary antibody (1:1000, BD Biosciences, 610397) in blocking solution containing 1.5% NGS, 0.1% Triton X100, 0.05% Tween-20 and 1% BSA. Following the primary antibody incubation, sections were rinsed in PBS and incubated in Alexa Fluor 647-conjugated goat anti-mouse (1:100, Invitrogen, A21235) secondary antibody in blocking solution containing 1.5% NGS, 0.1% Triton X100, 0.05% Tween 20 and 1% BSA for 2 hours at room temperature and protected from light. Sections were rinsed in PBS, mounted on positively charged glass slides, and left overnight to air-dry before coverslips were set using Fluoromount-G (SouthernBiotech). Representative high magnification images were collected using a Nikon A1R laser scanning confocal microscope and a 40x oil-immersion objective. Laser intensity, gain, and pinhole were kept constant between images. Sequential acquisition of multiple channels was used, and z-stacks were collected at 0.9 μm steps. Images were collected at a size of 0.7 x 0.59 mm and were automatically stitched upon acquisition using NIS Elements software. Image stacks were converted into maximum intensity z-projections using the NIS Elements software. Anatomical limits of each region were identified using a mouse brain atlas (Paxinos and Franklin, 2008).

### Sciatic nerve surgeries and nociceptive testing

#### Sciatic cuff implantation

Sciatic nerve cuff and sham surgeries were performed as previously described (Benbouzid et al., 2008). In brief, mice were anesthetized using 2% isoflurane (0.5 L/min). A 1-cm long incision was made along the proximal third of the lateral left thigh and the sciatic nerve was exteriorized and gently stretched using forceps. The nerve was either returned to its normal position (sham animals) or a 2 mm piece of PE-20 non-toxic, sterile polyethylene tubing (0.38 mm ID/1.09 mm OD) was split along and slid onto the sciatic nerve (cuff animals). After returning the nerve to its normal position, wound clips were used to close the skin. All electrophysiological and behavioral experiments were performed 6-14 days following surgeries.

#### Nociceptive testing

Mice were habituated to the 11 × 11 × 13 cm ventilated Plexiglas testing chambers placed on an elevated mesh platform (for von Frey and acetone tests) or a clear glass surface heated to 30°C (for Hargreaves test) for 1-3 h prior to testing. von Frey filaments (North Coast Medical, Inc. San Jose, CA) were used to assess mechanical sensitivity as previously described (Carrasquillo and Gereau IV, 2007). Beginning with the smallest fiber, the experimenter applied filaments to the hind-paw until the filament bent ∼ 30° for ∼2 s. The smallest filament to evoke a paw withdrawal response in at least three of five trials was recorded as the paw withdrawal threshold. Thermal sensitivity to heat was assessed using a modified version of the Hargreaves test (Hargreaves et al., 1988) as described previously (Carrasquillo and Gereau IV, 2007). A thermal stimulus with an active intensity of 35 was delivered from a constant radiant heat source through the glass bottom of the chamber to the plantar surface of the hind-paw (IITC Life Sciences, Woodland Hills, CA) and the latency to elicit paw withdrawal was recorded. To measure sensitivity to cold, we adapted the acetone evaporative test (Choi et al., 1994). Acetone (Sigma) was drawn into a 1 mL syringe and a drop was lightly applied to the hind-paw through the wire mesh. The drop of acetone (and not the syringe tip) was placed against the plantar surface of the hind-paw. Nociceptive responses and pain-like behaviors were quantified for 60 s following acetone exposure. Response quantification was modeled after a previously reported scoring system (Colburn et al., 2007). Transient lifting, licking, or shaking of the hind-paw that subsided immediately was given a score of 0; the same behaviors continuing up to but not past 5 s after initial application was given a score of 1; repeated and prolonged lifting, licking, or shaking of the hind-paw was given a score of 2. Three to five measurements were taken for each hind-paw on each behavioral assay and the average paw withdrawal thresholds (von Frey), paw withdrawal latencies (Hargreaves) and nociceptive scores (acetone) were calculated individually for each hindpaw. Hypersensitivity was assessed by comparing withdrawal thresholds in the paw ipsilateral to the side of sciatic nerve surgery compared to the paw contralateral to the side of sciatic nerve surgery.

### Statistics

Results are expressed as mean ± standard errors of the mean (SEM). Analysis was performed using either Student’s unpaired t-tests (with or without Welch’s correction for variance), Mann-Whitney U tests, Chi-squared (one-sided) tests, or two-way analyses of variance (ANOVA) followed by posthoc Tukey, Sidak’s or Dunnett’s multiple comparison tests. All analyses were performed using GraphPad Prism (v. 8) and *p* values lower than 0.05 were considered significant and are reported in figure legends. Detailed information for all statistical tests performed are reported in **Table 3**.

**Table 3.**
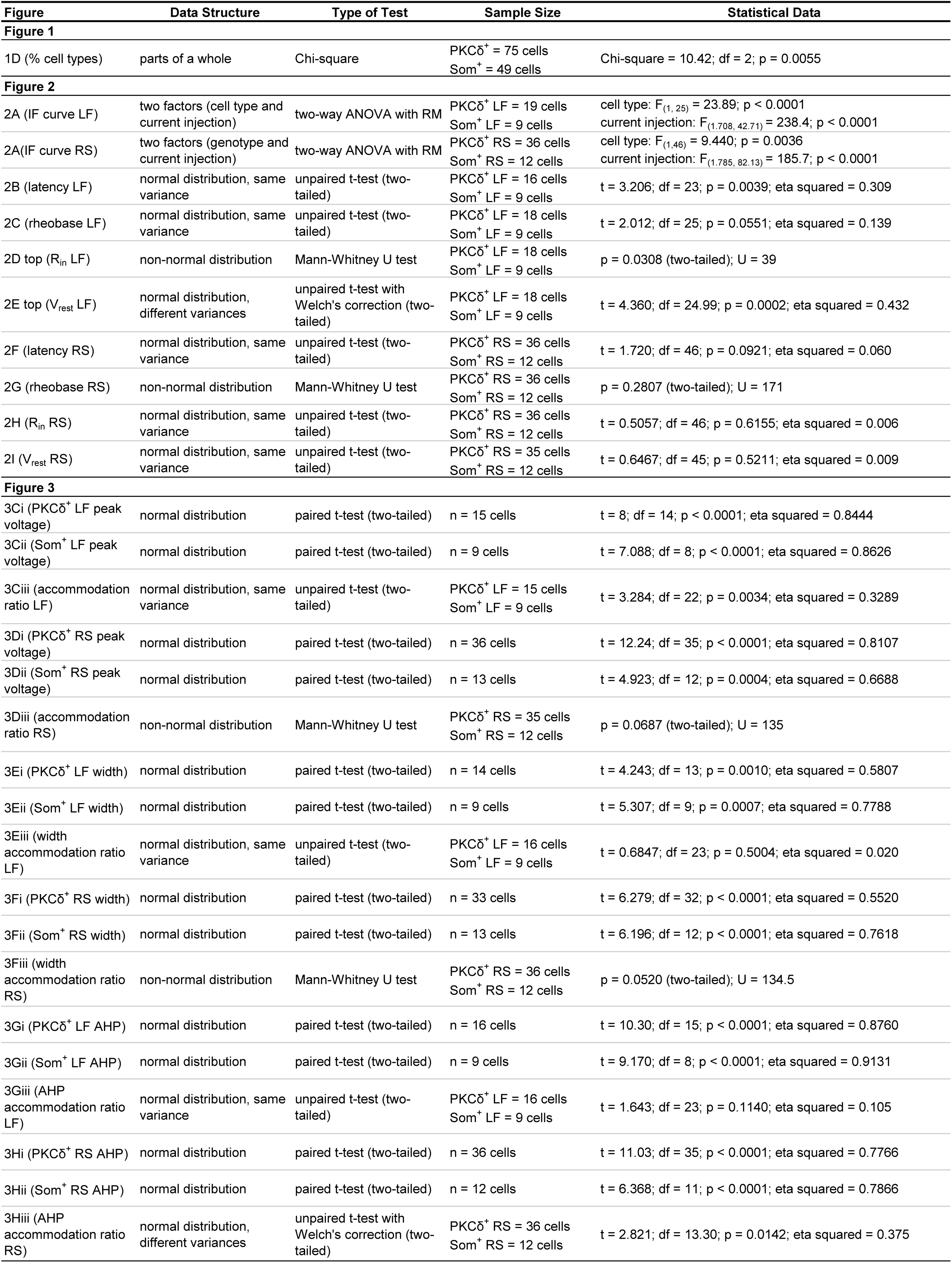
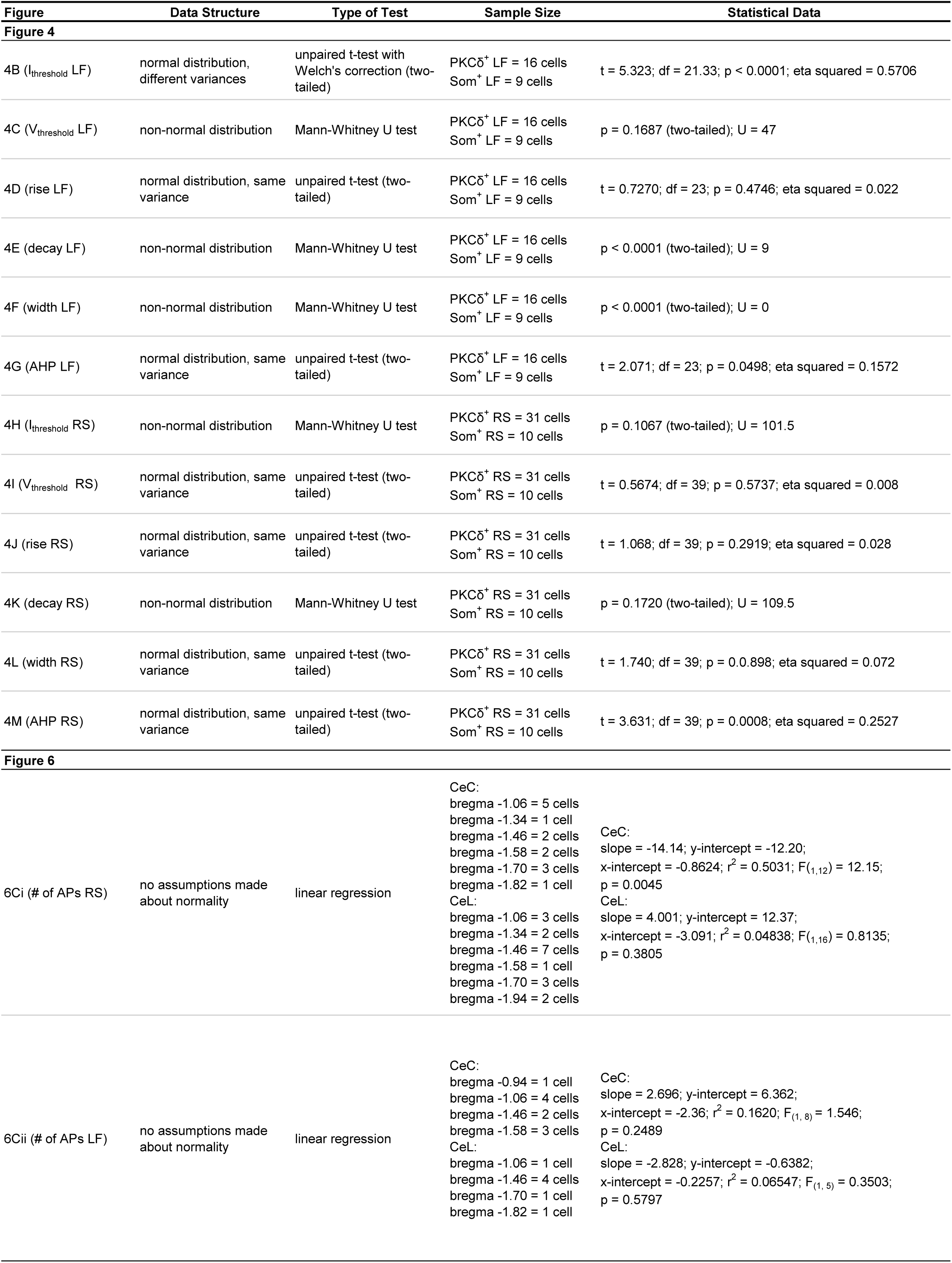
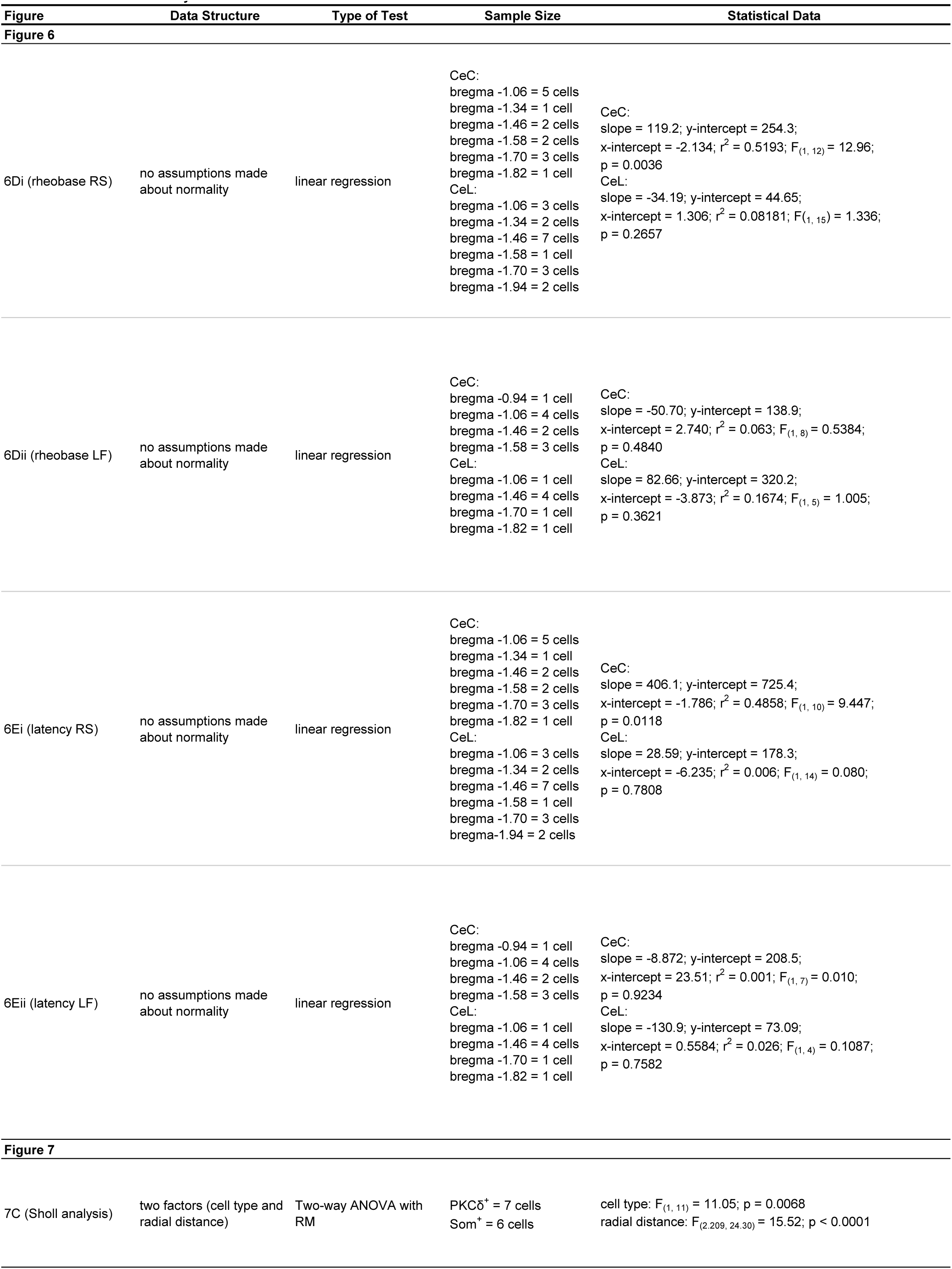
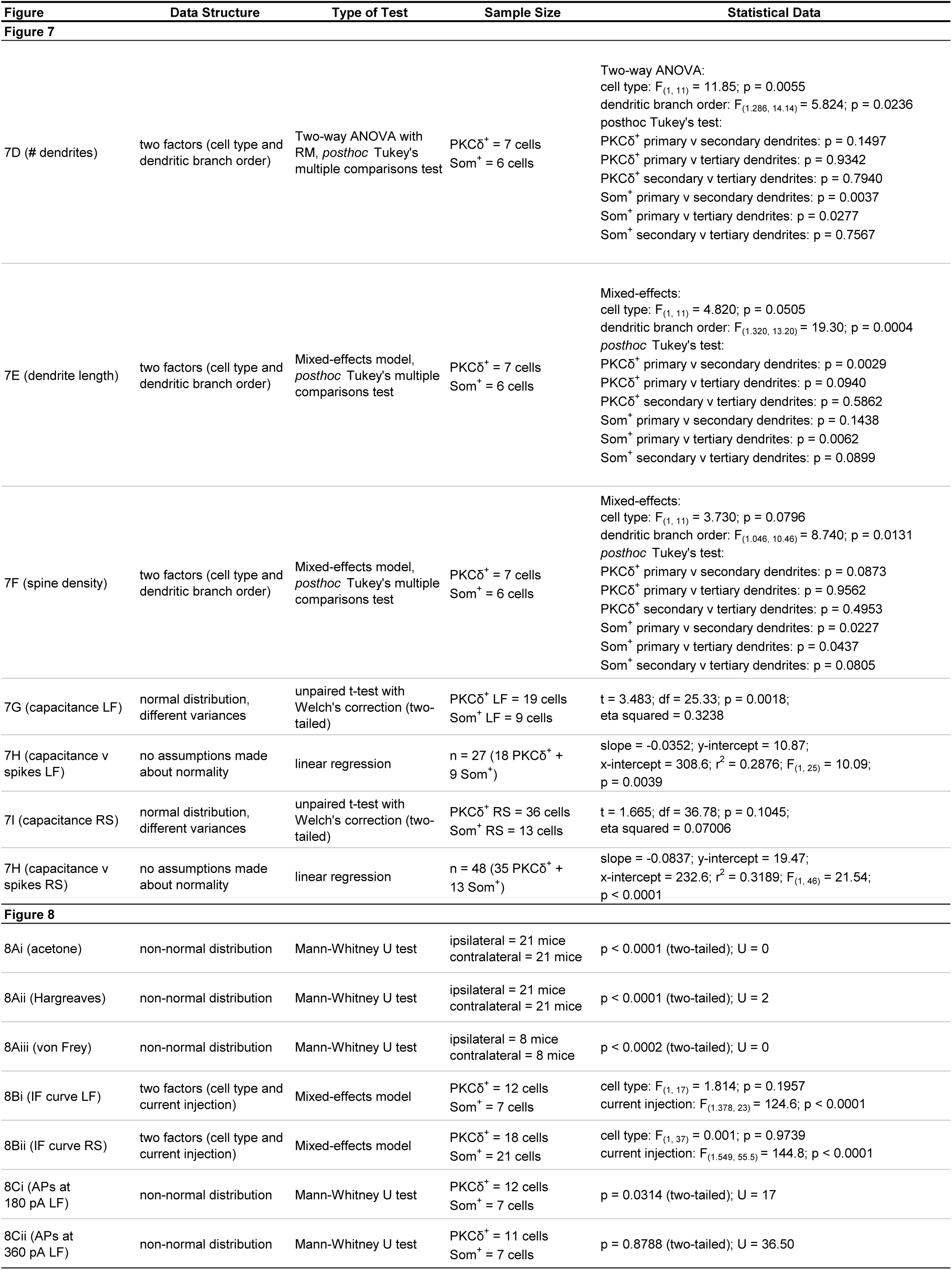
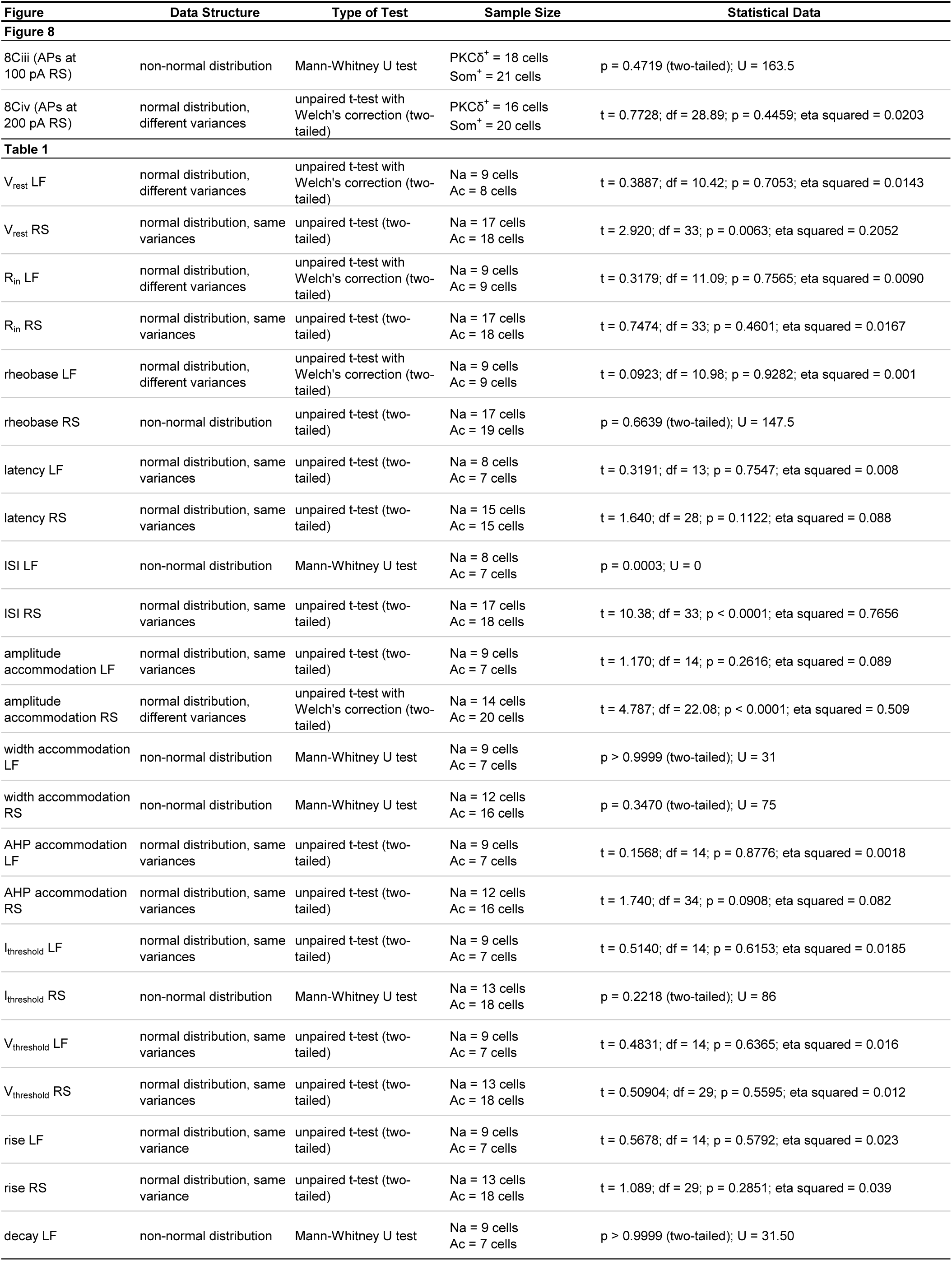
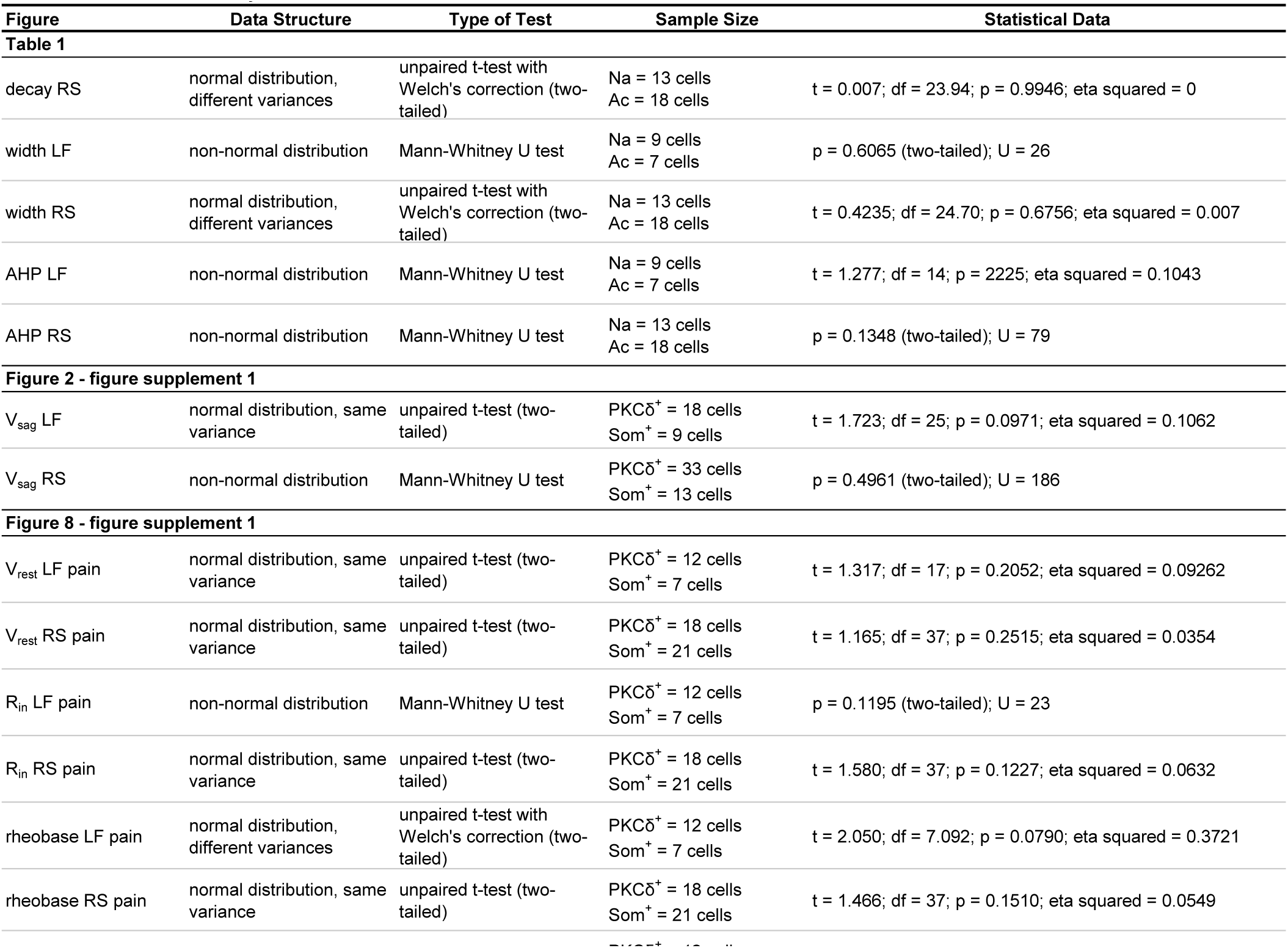
Statistical Analyses. Detailed information about data structure, statistical tests and results, and sample sizes. LF: late-firing; RS: regular spiking; IF: current-frequency plot; RM: repeated measures; F_(DFn, DFd)_: degree of freedom for the numerator of the F ratio, for the denominator of the F ratio; df: degrees of freedom; R_in_: input resistance; V_rest_: resting membrane potential; AHP: afterhyperpolarization; I_threshold_: current threshold; F ratio; df: degrees of freedom; R_in_: input resistance; V_rest_: resting membrane potential; AHP: afterhyperpolarization; I_threshold_: current threshold; V_threshold_: voltage threshold; AP: action potential; Na: non-accommodating; Ac: accommodating; ISI: interspike interval; V_sag_: voltage sag.

### Data availability

All data in this study is available from the corresponding author.

## ACKNOWLEDGEMENTS

This research was supported by the National Center for Complementary and Integrative Health Intramural Research Program. We would like to thank Drs. Hugo Tejeda, Mario Penzo and Yavin Shaham for comments on this manuscript and Dr. Ted Usdin and the Systems Neuroscience Imaging Resource of the National Institute of Mental Health for making possible the morphological reconstruction and analysis of biocytin-filled cells. We would also like to thank the National Institutes of Neurological Disorder Mouse Facility staff for their vital work in animal husbandry.

## AUTHOR CONTRIBUTIONS

Conceptualization, Y.C.; Methodology, Y.C., A.P.A. and S.V.; Investigation, A.P.A., A. K., H.-S.A., J. J. B., T.D.W., S.V., Y. K. S, and S. M. G; Writing, Y.C. and A.P.A.; Supervision, Y. C.; Funding Acquisition, Y.C.

---

**Figure 1 – figure supplement 1.**
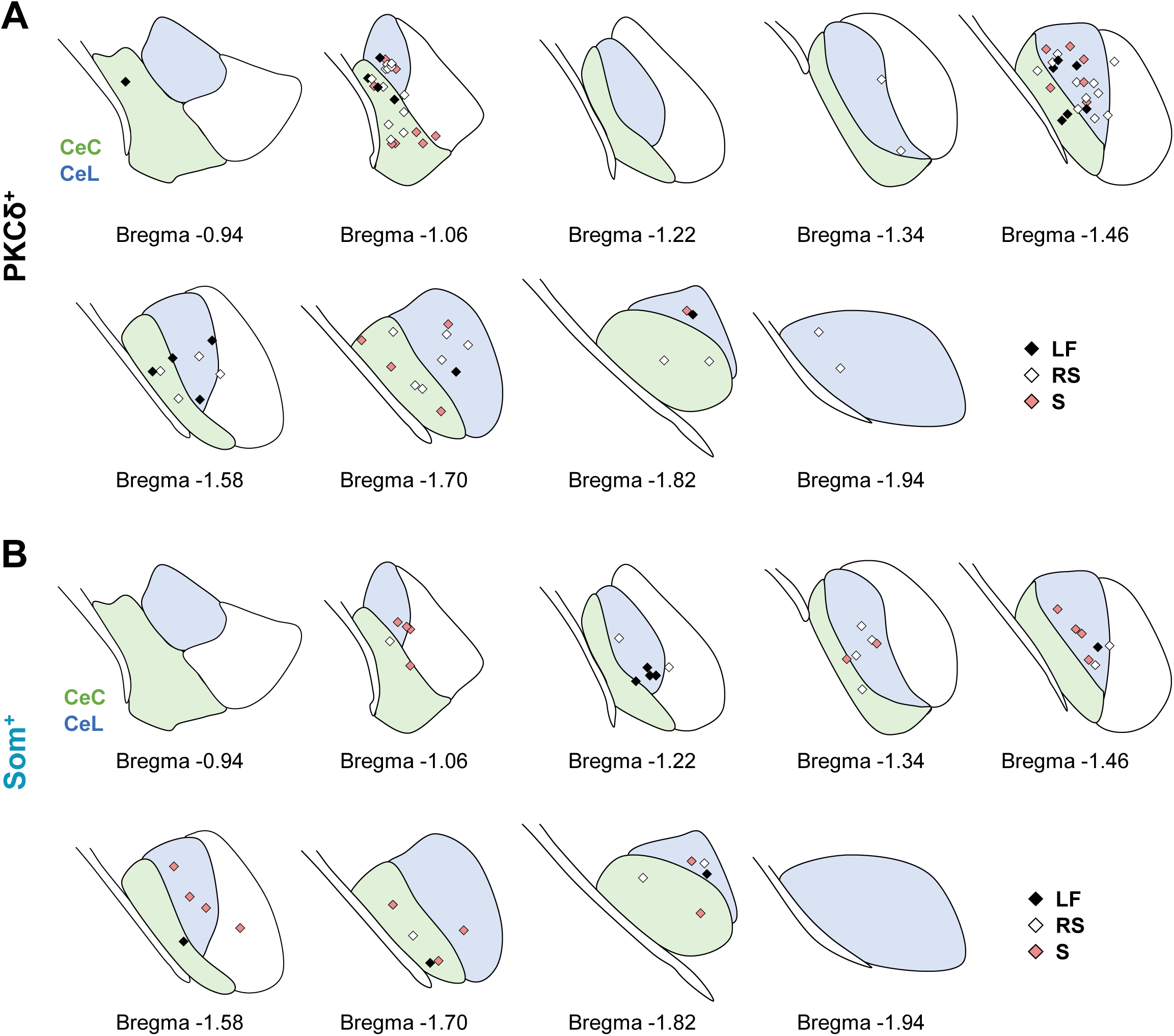
Anatomical location of electrophysiology recordings. Rostro-caudal anatomical locations of recorded PKCδ^+^ (**A**) and Som^+^ (**B**) cells, represented as a schematic of the CeLC, created using Paxinos & Franklin, 2008. The capsular (CeC) and lateral (CeL) subdivisions of the central amygdala are shown in green and blue, respectively. LF = late-firing; RS = regular-spiking; S = spontaneous.

**Figure 2 – figure supplement 1.**
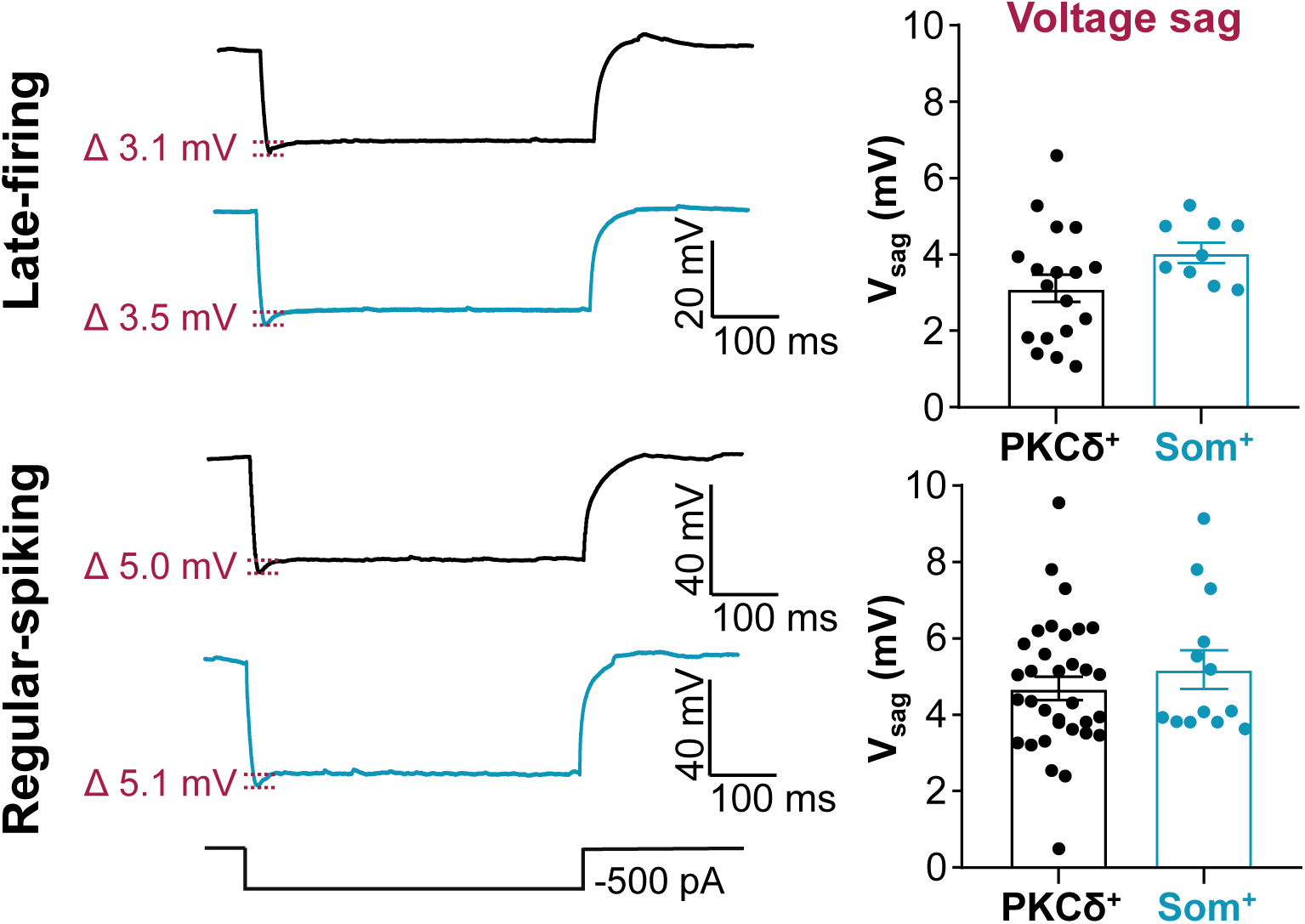
Voltage sag is indistinguishable in PKCδ^+^ and Som^+^ cells. Representative traces of late-firing (top) and regular-spiking (bottom) neurons in response to a 500 ms hyperpolarizing current injection, with PKCδ^+^ cells shown in black and Som^+^ cells in blue. Values are reported as mean ± S.E.M. For PKCδ^+^ cells: n = 18 cells for late-firing and n = 33 regular-spiking. For Som^+^ cells: n = 9 for late-firing and n = 13 for regular-spiking. Voltage sag = V_sag_

**Figure 8 – figure supplement 1.**
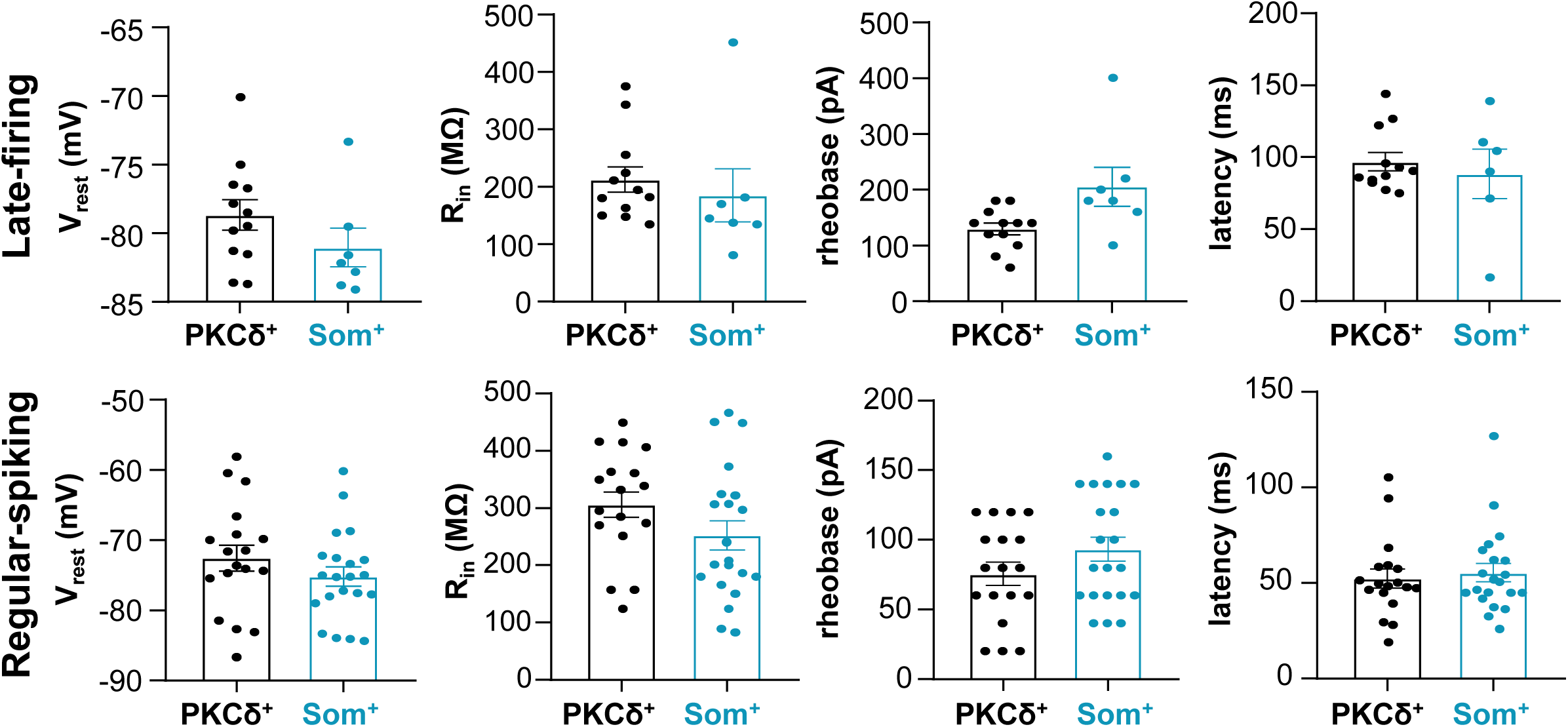
Passive membrane and repetitive firing properties of PKCδ^+^ and Som^+^ cells following nerve injury. All data are reported as mean ± S.E.M. For PKCδ^+^ cells: n = 12 cells for late-firing and n = 18 regular-spiking. For Som^+^ cells: n = 6-7 for late-firing and n = 21 for regular-spiking. V_rest_ = resting membrane potential; R_in_ = input resistance.

